# Comprehensive analysis of hsa-miR-654-5p’s tumor-suppressing functions

**DOI:** 10.1101/2020.12.20.423719

**Authors:** Chuanyang Liu, Lu Min, Jingyu Kuang, Chu-shu Zhu, Xinyuan Qiu, Xiaomin Wu, Tianyi Zhang, Sisi Xie, Lingyun Zhu

**Affiliations:** Department of Biology and Chemistry, College of Liberal Arts and Sciences, National University of Defense Technology, Changsha, Hunan, 410073, China

**Keywords:** pan-cancer, bioinformatics, hub target genes, miR-654-5p, epithelial-mesenchymal transition

## Abstract

The pivotal roles of miRNAs in carcinogenesis, metastasis, and prognosis have been demonstrated recently in various cancers. This study intended to investigate the specific roles of hsa-miR-654-5p in lung cancer, which was poorly discussed. A series of closed-loop bioinformatic functional analyses were integrated with *in vitro* experimental validation to explore the overall biological functions and pan-cancer regulation pattern of miR-654-5p. We found that miR-654-5p abundance was significantly elevated in LUAD tissues and correlated with patients’ survival. 275 potential targets of miR-654-5p were then identified and the miR-654-5p-RNF8 regulation axis was validated *in vitro* as a proof of concept. Furthermore, we revealed the tumor-suppressing roles of miR-654-5p and demonstrated that miR-654-5p inhibited lung cancer cell epithelial-mesenchymal transition (EMT) process, cell proliferation, and migration using target-based, abundance-based and ssGSEA-based bioinformatic methods and *in vitro* validation. Following the construction of a protein-protein interaction network, 11 highly interconnected hub genes were identified and a five genes risk scoring model was developed to assess their potential prognostic ability. Our study will not only provide a basic miRNA-mRNA-phenotypes reference map for understanding the function of miR-654-5p in different cancers but also reveal the tumor-suppressing roles and prognostic values of miR-654-5p in lung cancer.

**Graphical abstract:** 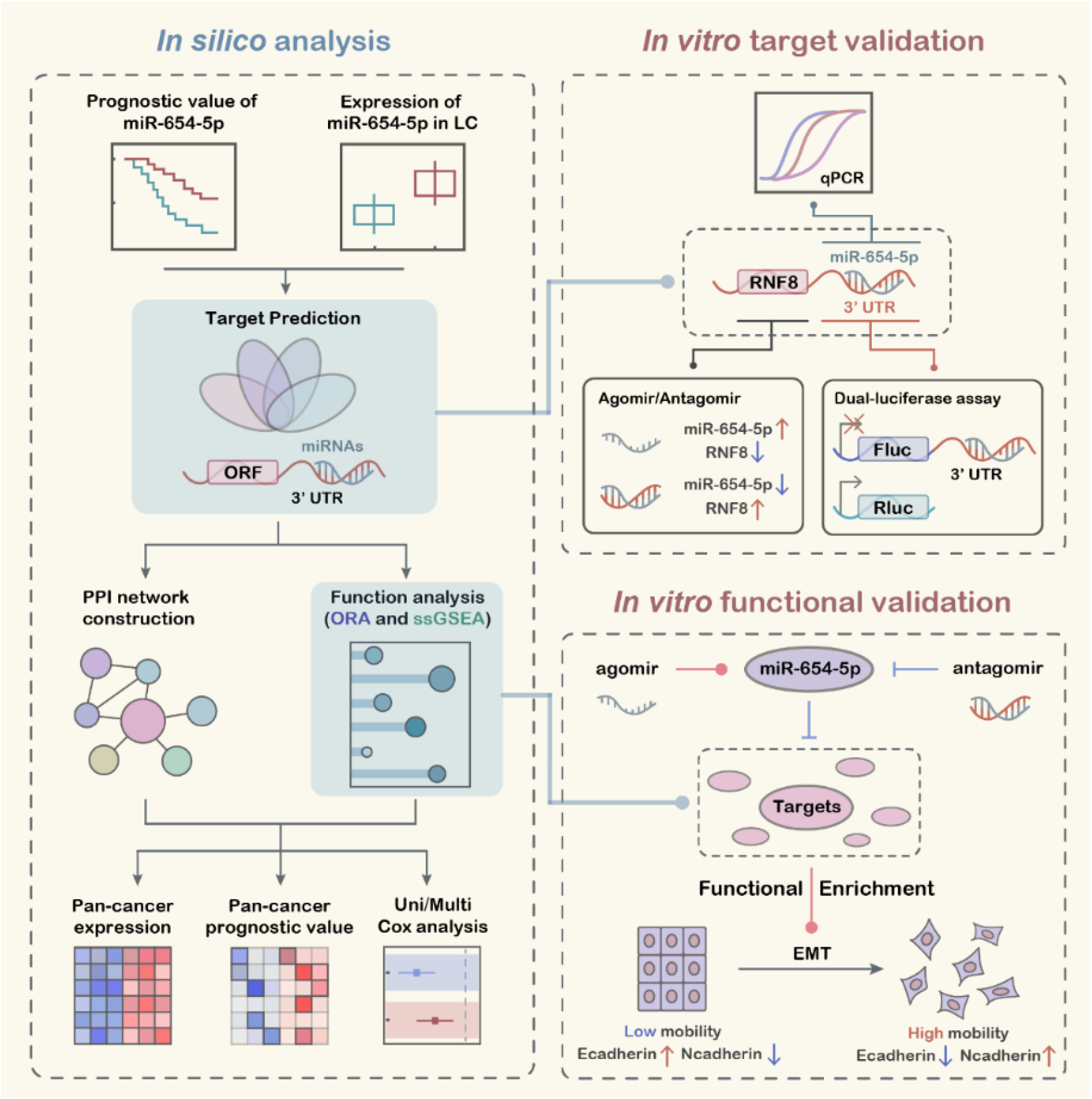

## 1 INTRODUCTION

Lung cancer is one of the most malignant types of cancer and constitutes the primary cause of cancer-related deaths worldwide^1-4^. Non-small cell lung cancer (NSCLC), the dominant type of lung cancer with high mortality, accounts for approximately 80% of all lung cancer cases. The main cause of its high mortality is that cancer cells can easily metastasize to vital organs and finally lead to multiple organ failure^5-7^. As a consequence, survival for small cell lung cancer remained low worldwide, highlighting the importance of mechanistic research about NSCLC and metastasis^4,8^. However, the detailed molecular mechanism of non-small cell lung cancer metastasis remains unclear at present, which largely limits the development of target-driven drugs and therapeutics.

Recently, a class of short non-coding RNAs, microRNAs (miRNAs), have been proven to play important roles in cancer metastasis via post-transcriptional regulation of their target genes’ expression^9-12^. For example, downregulation of has-miR-654-3p (miR-654-5p) in colorectal cancer promotes cell proliferation and invasion by targeting SRC^13^. Xiong et al revealed that overexpression of miR-654-3p suppresses cell viability and promotes apoptosis by targeting RASAL2 in non-small-cell lung cancer. In addition, hsa-miR-654-5p (miR-654-5p), derived from the same precursor miRNA as miR-654-3p, showed different biological functions in cancers. miR-654-5p has been reported to attenuate breast cancer progression by targeting EPSTI1^14^. In osteosarcoma, miR-654-5p inhibits tumor growth by targeting SIRT6^15^. In addition, miR-654-5p targets GRAP to promote proliferation, metastasis, and chemoresistance of oral squamous cell carcinoma through Ras/MAPK signaling^16^. These studies provide some specific explanations for how miR-654-5p regulates certain biological processes in cancers. However, in a larger landscape, the overall functions of miR-654-5p and how it regulates cellular pathway networks or biological processes have been poorly investigated and assessed, and the precise role of miR-654-5p in lung cancer remains poorly discussed except for Kong’s work, which demonstrated that Hsa_circ_0085131 acted as a competing endogenous RNA of miR-654-5p to release autophagy-associated factor ATG7 expression and promoted cell chemoresistance^17^. Thus, the identification and comprehensive functional analysis of miR-654-5p target genes would provide more insight into the mechanism underlying miR-654-5p-induced downstream signaling transduction or phenotype alterations in cancers.

Here, we designed a series of experiments integrating bioinformatics analysis and *in vitro* validation experiments, aiming to elucidate the main functions and regulation pattern of miR-654-5p from all angles.

## 2 MATERIALS AND METHODS

### 2.1 Pan-cancer analysis of hsa-miR-654-5p

Pan-cancer overall survival and expression analysis were performed based on data downloaded from UCSC Xena Pan-Cancer Atlas Hub (https://xenabrowser.net/hub/)^18^. Lung cancer tissue samples data (GSE169587, including 12 normal tissues and 38 lung cancer tissues) and serum data (GSE152072, including 40 LUAD samples, 38 LUSC samples and 61 healthy/non-diseased samples) were downloaded from NCBI GEO database (https://www.ncbi.nlm.nih.gov/geo/). The R package “survival” and “survminer” was used for survival analysis and visualization of Kaplan-meier survival analysis. Optimal cutoff of hsa-miR-654-5p expression was obtained using function “surv_cutpoint” ^19^.

### 2.2 Integrated target prediction of miR-654-5p target genes

We analyzed the potential binding between miR-654-5p and potential target genes using miRWalks (http://mirwalk.umm.uni-heidelberg.de/)^20^ and 3 other highly recognizable miRNA-target prediction tools (miRanda: http://www.microrna.org/microrna/getMirnaForm.do, miRDB: http://mirdb.org/ and Targetscan7.2: http://www.targetscan.org/vert_72/). The results were analyzed and visualized by R package “VennDiagram” and Cytoscape software (version 3.8.2).

### 2.3 Comprehensive analysis of miR-654-5p functions

#### 2.3.1 Functional analysis of hsa-miR-654-5p

The 275 overlapping gene symbols were converted to Entrez ID and submitted to clusterProfiler, a user-friendly R package designed for statistical analysis and visualization of functional profiles for gene clusters^21^, and the Metascape (metascape.org/gp/index.html)^22^. Gene ontology (GO) annotation, KEGG pathway enrichment analysis and other over representation analyses (ORA) were performed and adjusted p-value<0.05 was considered as significant. Furthermore, Hallmark (p<0.05), KEGG functional sets (p<0.05), Oncogenic signatures (p<0.05), Reactome (p<0.001), BioCarta (p<0.05) and Canonical pathway (p<0.01) enrichment analysis enrichment analysis were also performed by KOBAS database. In addition, the predicted targets of miR-654-5p by miRWalks 3.0 (with score>0.95) were also submit to its own GO annotation and KEGG pathway enrichment analysis to avoid missing key information after only considering the intersection of predicted targets. R package “ggplot2” and “enrichplot” were used to visualized the results of ORA analysis. The ssGSEA (single sample gene set enrichment analysis) scores for 1387 constituent pathways in NCI-PID, BioCarta and Reactome were calculated by the PARADIGM algorithm. The correlation between hsa-miR-654-5p and EMT score in LUAD and LUSC were visualized by R package “ggpubr” and a p-value<0.05 was considered as significant.

#### 2.3.2 Protein-protein interaction analysis

Protein-protein interaction (PPI) network of overlapping genes was construct by the Search Tool for the Retrieval of Interacting Genes (STRING, https://string-db.org/). High confidence (minimum required interaction score>0.700) was set as the selection criterion of constructing the network. The PPI was downloaded for further analysis and visualized by Cytoscape software (version 3.8.2). Molecular Complex Detection (MCODE) plugin in Cytoscape was utilized to find potential modules in the PPI network based on topology. The degree cut-off value to 2 and the node score cut-off to 0.2 were set in the MCODE process. Genes with those differentially expressed in lung adenocarcinoma (LUAD) and highly connected (Degree >2) were selected as hub genes.

#### 2.3.3 Pan-cancer expression and prognostic value of hub genes

Pan-cancer Batch effects normalized mRNA data (n=11,060) were downloaded from UCSC Xena Pan-Cancer Atlas Hub. The pan-cancer expression of hub genes was shown as a heatmap using the R package “pheatmap”, The non-parametric test (Wilcoxon rank sum test) was used to compare the means of gene expression. The R package “survival” and “survminer” was used for Kaplan-Meier survival analysis^19^. To further assess the prognostic value of these genes, Gene Expression Profiling Interactive Analysis (GEPIA) tool (http://gepia.cancer-pku.cn/), including integrated TCGA mRNA sequencing data and the GTEx, were also used (with FDR P-value adjustment, 0.05 significance level, and Median group cut-off) to calculate patient overall survival rate (OS) and relapse-free survival rate (RFS) ^23,24^.

#### 2.3.4 hsa-miR-654-5p expression-based functional analysis

Pan-cancer Batch effects normalized mRNA data set and miRNA data set were used. Only LUAD samples were chosen for further analysis. Cancer samples were divided into three groups based on the abundance of hsa-miR-654-5p. R package “limma” was used to perform differential expression analysis between high abundance group and low abundance group. DEGs were obtained with an automatically generated criterion (mean(abs(logFC)) + 2*sd(abs(logFC)) and adj. p-value < 0.05). The r package “clusterProfiler”, “DOSE”, “msigdbr” and “enrichplot” were used for enrichment analysis of DEGs list and subsequent visualization. ssGSEA scores of TCGA samples were downloaded from UCSC Xena Pan-Cancer Atlas Hub, visualization of ssGSEA analysis were performed using “ggplot2” and “ggpubr”.

#### 2.3.5 Cox model construction and validation

Pan-cancer Batch effects normalized LUAD mRNA data set (n=514) was randomly divided into two cohort, training cohort (n=308) and testing cohort (n=206). 11 hub genes were chosen and univariate/multivariate Cox regression was performed to evaluate the prognostic value of hub genes. R package “survival” and function “coxph” was used. A risk model based on the expression of genes and coefficients in cox regression analysis was constructed. The risk score of each cancer sample was calculated as:

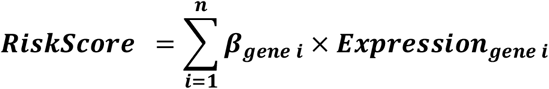

Based on the value of *RiskScore*, cancer samples in the Pan-cancer data hub are divided into the high-risk group and the low-risk group. And optimal cutoff of *Riskscore* was automatically generated using the function “surv_cutpoint”. A log-rank p<0.05 was considered significant^19^. The ROC (Receiver Operating Characteristic) curve was calculated using R package “pROC”^25^.

### 2.4 *In vitro* proof of concept for functional analyses

#### 2.4.1 Cell culture

Lung cancer cell lines A549, H1299 and human embryonic kidney cell line 293T were purchased from American Type Culture Collection (ATCC, Manassas, VA), and cultured in RPMI 1640 Medium (Hyclone) supplemented with 10% fetal bovine serum (FBS; GIBCO, Gaithersburg, MD, USA) and 100 U/ml penicillin and streptomycin (P/S; Hyclone). Cells were contained in a 5% CO_2_ incubator at 37°C.

#### 2.4.2 Overexpression or knockdown of miRNA and RNF8

*in vitro* overexpression and knockdown of miR-654-5p were performed by Lipofectamine^™^ 3000 (Invitrogen)-mediated miRNA Agomir and Antagomir transfection. Chemically modified hsa-miR-654-5p mimics and hsa-miR-654-5p inhibitor, with their negative control Stable N.C and inhibitor N.C respectively, were purchased from GenePharma. A549, H1299 and HEK293T cells were transfected according to the manufacturer’s directions with the working concentration of Antagomir/Agomir 20nM.

#### 2.4.3 RNA extraction and quantitative real-time PCR

Total RNAs were extracted using the TRIzol agent (Ambion) according to the instruction of the manufacturer. Reverse transcription of RNA and quantitative real-time PCR was performed using the Hairpin-it^™^ miRNAs qPCR Quantitation Assay Kit (GenePharma) according to the manufacturer’s instructions. Quantitative RT-PCR was performed in a Roche 480 real-time PCR system. The 2^-ΔΔCt^ method was used to evaluate the miR-654-5p gene expression after normalization for expression of the endogenous controls U6 (U6 non-coding small nuclear RNA). All primers for miR-654-5p and the U6 genes were synthesized and approved by GenePharma. Each experiment was repeated at least three times.

#### 2.4.4 Western Blotting

Total proteins of cells were extracted using RIPA lysis buffer (Beyotime) with Protease Inhibitor (Roche) and Phosphatase Inhibitor (Roche). Protein samples were separated in sodium dodecyl sulfate (SDS)-PAGE and transferred to polyvinylidene fluoride (PVDF) filter membranes (Millipore, USA) for immune-hybridization. After 1 hour blocking in PBST (phosphate buffered saline containing 0.05% Tween-20 and 5% non-fat milk powder), the membranes were incubated with one of the following primary antibodies with corresponding concentration: RNF8 (Santa Cruz Biotech, 1:500), EMT kit (Cell Signaling Technology, 1:2000), beta-actin (Santa Cruz Biotech,1:4000), Secondary antibodies were Horseradish peroxidase (HRP)-conjugated anti-mouse IgG (ZB-2305,ZSGB-Bio,1:4000) or anti-Rabbit IgG (ZB-2301, ZSGB-Bio, 1:4000). Subsequently, band visualization was performed by electro-chemiluminescence (ECL) and detected by Digit imaging system (Thermo, Japan).

#### 2.4.5 *In vitro* cell proliferation and wound healing assays

To perform cell proliferation assays, cells were counted and plated in the well of 96-well plate (1500 cells per well) 24h after transfection of chemically modified oligonucleotides. the cell proliferation ability was determined using the Cell Counting Kit-8(CCK8) Assay Kit (Dojindo Corp, Japan) according to the manufacturer’s protocol. Cell migration assays were performed using a 6-well plate. 24h after transfection, cells should reach ∼70-80% confluence. 20μL pipette tip were utilized to scratch the monolayer cells after washing by PBS. Gently wash the well twice with PBS again to remove the detached cells and replenish the well with fresh complete medium. After 24 hours incubation. Cell migration was imaged by microscope and quantified by imageJ software. The assay was performed three independent times.

### 2.5 Statistical analysis

Statistical analysis was conducted using the GraphPad Prism (Version 8.0; SPSS Inc.). All results were presented as the mean ± standard error of the mean (SEM). Student *t*-test, Wilcoxon rank sum test was performed to compare the differences between treated groups relative to their controls. p-values are indicated in the text and figures above the two groups compared and p<0.05 (denoted by asterisks) was considered as statistically significant.

## 3 RESULTS

### 3.1 *In silico* prognostic analysis of miR-654-5p in cancers

To assess the prognostic value of miR-654-5p in various cancers, we performed a pan-cancer survival analysis based on the Pan-cancer TCGA data set^26^. The results showed that high abundance of miR-654-5p was significantly associated with poor prognosis of ACC (Fig. S1A, p = 5×10^−3^), BRCA (Fig 1A, p = 3.4×10^−3^), HNSC (Fig 1B, p = 1.6×10^−3^), KIRC (Fig. S1B, p = 5.3×10^−3^), LGG (Fig 1C, p = 1.8×10^−3^), LIHC (Fig. S1D, p = 6.5×10^−3^) and THCA (Fig 1D, p < 6.5×10^−3^). A similar pattern can be observed in BLCA, COAD, KICH, KIRP, PCPG, SKCM, STAD and THYM (Fig. S1). In LAML (Fig. S1C, p =4.4×10^−3^), MESO (Fig. S1F, p =7.5×10^−3^), as well as LUSC, PAAD and READ (Fig. S1), high abundance of hsa-miR-654-5p is significantly associated with better prognosis. These results indicated that miR-654-5p might be related to the progression of multiple cancers. In other cancers, however, the relation of hsa-miR-654-5p and survival rate were not significant. Pan-cancer expression analysis revealed that miR-654-5p abundance was significantly elevated in ESCA, LUAD, LUSC and STAD tumor tissues, while significant down-regulation of miR-654-5p was found in BRCA, COAD, HNSC, KICH, KIRP, LIHC, READ and THCA (Fig 1E), Strikingly, the role of miR-654-5p in STAD and LUSC seems to be contradictory. For lung cancer, hsa-miR-654-5p was significantly upregulated both in LUAD and LUSC samples compared to normal (Fig 1F and G). These results were further validated in external datasets (Fig 1H∼I). hsa-miR-654-5p is also elevated in the serum of lung cancer patients (Fig 1J). Taken together, these results indicated that hsa-miR-654-5p might be associated with the lung cancer tumorigenesis and progression.

**Figure 1.**
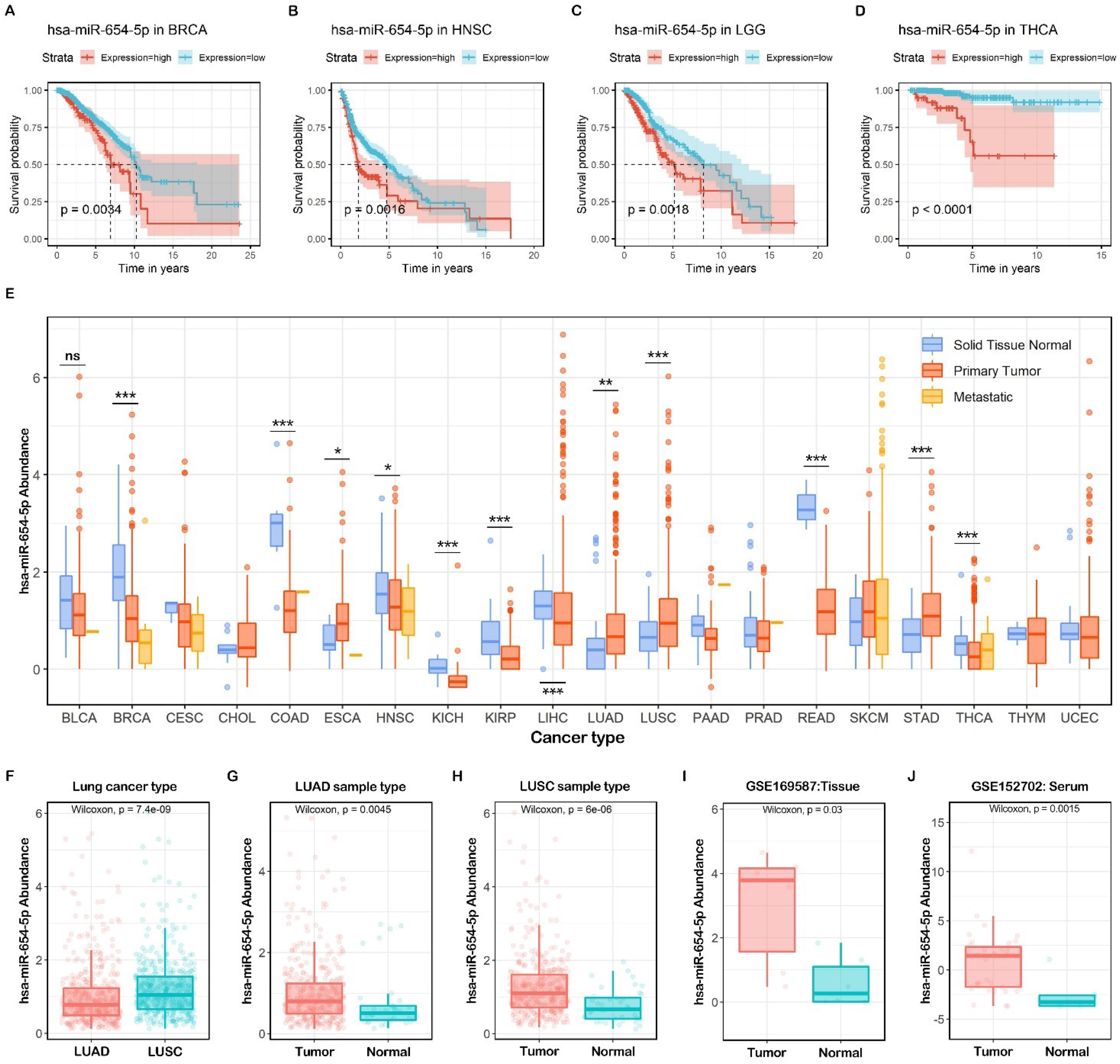
Pan-cancer expression pattern and survival analysis of hsa-miR-654-5p. (A-D) Overall survival (OS) of hsa-miR-654-5p based on TCGA pan-cancer normalized miRNA dataset. (E) hsa-miR-654-5p abundance in TCGA pan-cancer normalized miRNA dataset. (F∼I) the abundance of hsa-miR-654-5p in lung cancer tumor samples and normal tissues. (J) the abundance of hsa-miR-654-5p in the serum of lung cancer patients and healthy people.

### 3.2 Integrated targets prediction of miR-654-5p

The classic function of miRNAs is to bind to the 3’ untranslated region (3’UTR) and inhibit the translation of target gene mRNA^11,27,28^. To reveal the common biological functions of miR-654-5p, we utilized multiple bioinformatic tools, including miRWalks 3.0 and three other highly promising miRNA-target prediction tools, to identify its potential targets. The intersection of these four predicted targets’ sets was subsequently analyzed and visualized (Fig. 2 A)^29,30^. As shown in the results, 275 overlapping genes were predicted by all four tools (Fig. 2 B), indicating that these genes should be promising targets of hsa-miR-654-5p. Among these genes, RNF8, CYP4A11, and WASF2 showed high target site accessibility, as shown in Fig 2B.

**Figure 2.**
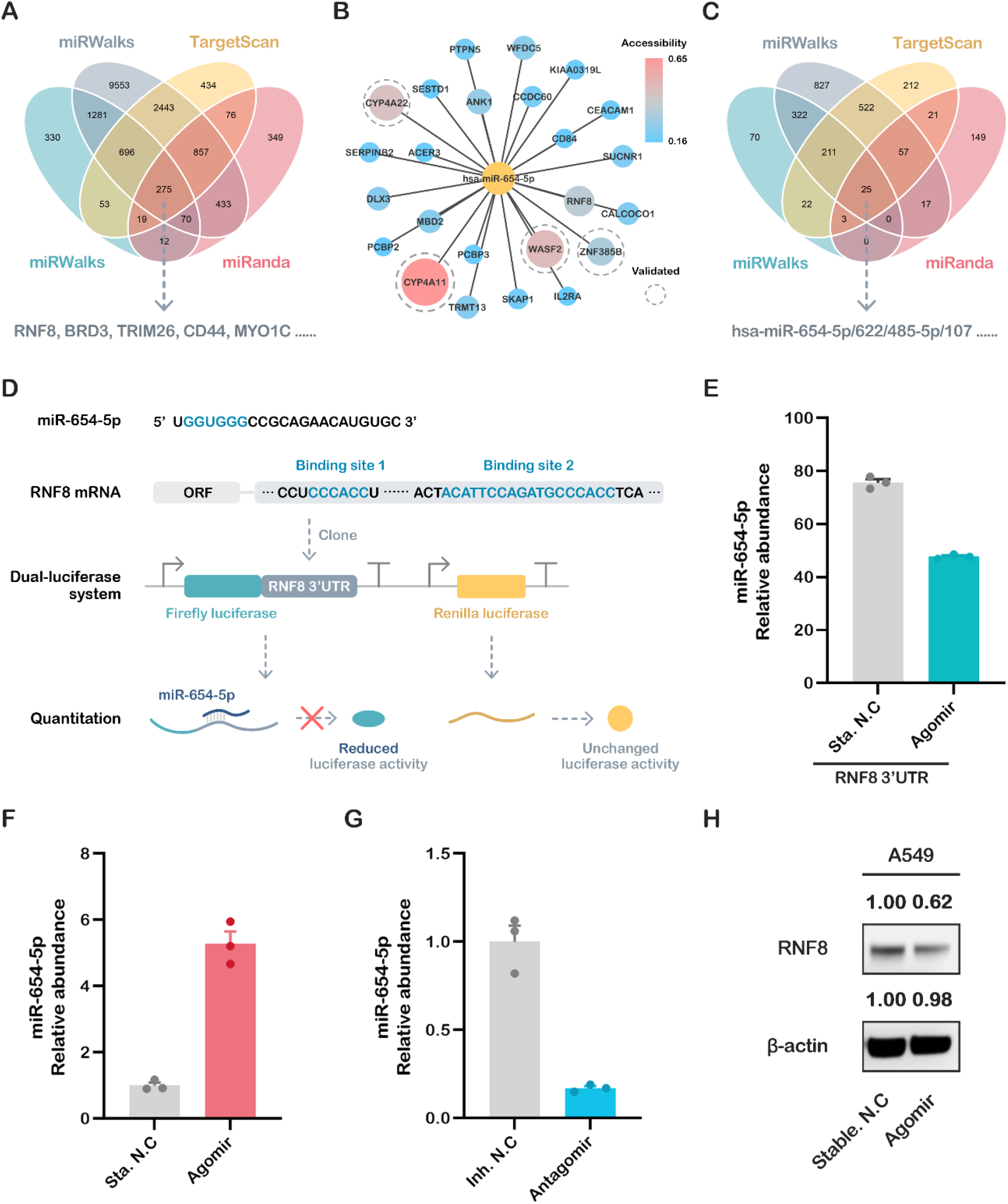
Prediction and validation of hsa-miR-654-5p target genes. (A) Four promising miRNA-targets prediction tools including miRanda, TargetScan7.2, RNA22 v2.0, and miRWalks v3.0 were used to find the targets of miR-654-5p. (B) miR-654-5p-targets interaction network was visualized using Cytoscape 3.8.2, the deeper color and larger sizes of nodes indicate the high accessibility of each gene. (C) Four tools were utilized to reverse miRNAs prediction using RNF8 3’UTR sequencing. (D) the binding between miR-654-5p and RNF8 3’UTR and the design of dual-luciferase plasmid. (E) 48 hours after the liposome-mediated transfection of miR-654-5p agomir/stable NC and designed dual luciferase plasmid, the dual-luciferase assay was performed. Relative light unit (RLU) was obtained and the ratio of Firefly luciferase activity/Renilla luciferase activity was calculated to evaluate the binding of miRNA and its target. (F and G) Real-time PCR was performed to detect the abundance of miR-654-5p in A549 cells treated with agomir/antagomir. (H) Western blot assay was performed to measure the expression of RNF8 in miR-654-5p Agomir treated cells. miR-654-5p RNA levels are expressed as the mean ± SEM of four different experiments normalized to U6 abundance. *, p<0.05, **, p<0.01, ***p<0.001 vs. control.

### 3.3 miR-654-5p/RNF8 axis: a case of *in vitro* validation of predicted targets

RNF8, a predicted target hub gene that has a high target site accessibility, has recently been proven to have a strong relation to breast cancer and lung cancer^31-34^. In addition, reverse miRNAs prediction using RNF8 3’UTR also indicated that miR-654-5p is one of the 25 miRNAs predicted by all four tools (Fig. 2C), which further implied that the axis of miR-654-5p-RNF8 might be pivotal to the biological roles of both miR-654-5p and RNF8 in cell regulation network. We therefore chose RNF8 as an *in vitro* proof of concept.

To validate the direct regulation of miR-654-5p on RNF8, the potential binding sites of miR-654-5p in the RNF8 3’UTR were screened, and two potential binding sites were found (Fig. 2D). We cloned the full-length 3’UTR sequence of RNF8 linked it to the 3’ end of the coding sequence of luciferase to mimic the natural transcriptional inhibition of miR-654-5p on RNF8. The dual-luciferase assay was then performed, and the results showed that the activity of luciferase was decreased significantly in the miR-654-5p-transfected group compared to the control (Fig. 2E, p <0.001), indicating that miR-654-5p could bind to 3’UTR and inhibit the transcription of RNF8.

To further prove that miR-654-5p inhibits the expression of RNF8 *in vitro*, lung cancer cells A549 were transfected with miR-654-5p agomir or antagomir. The results showed that miR-654-5p’s abundance in A549 was increased by 5.23-fold (Fig. 2F, p<0.001) compared to the control group following the agomir-transfection, and the RNF8 protein level was dramatically downregulated accordingly (Fig. 2G). Taken together, these results indicate that miR-654-5p reduces RNF8 expression via post-transcriptional inhibition, which is consistent with our previous prediction.

### 3.4 Comprehensive analysis of the predicted target genes and *in vitro* validation

#### 3.4.1 Functional enrichment analyses reveal the inhibitory roles of miR-654-5p on lung cancer cell proliferation

To obtain a better understanding of regulation pattern and core functions of miR-654-5p at a cellular level, over representation analysis (ORA) including GO annotation, and enrichment analyses were performed to the predicted target gene list of miR-654-5p. As shown in the results, molecular functions (MFs) terms such as kinase activity and growth factor receptor binding were significantly enriched (Fig. 3A). In GO biological processes (BPs) enrichment, these 275 targets were mainly enriched by the RTK signaling pathway and cellular response to growth factor stimulus. Multiple cell adhesion-related items such as regulation of cell-cell adhesion were also enriched, demonstrating miR-654-5p is likely to regulate cell adhesion (Fig. 3C). For cellular components (CCs), target genes were commonly enriched by cytoplasmic side of plasma membrane, cytoplasmic side of membrane (Fig. 3B). Furthermore, RTK pathway-related pathways such as VEGF and Ras signaling were enriched in KEGG pathway enrichment, consistent with GO BPs results (Fig. 3D). Taken together, these results indicate miR-654-5p might be crucial in cell adhesion processes and RTK-related biological processes such as growth factor-regulated cell proliferation.

**Figure 3.**
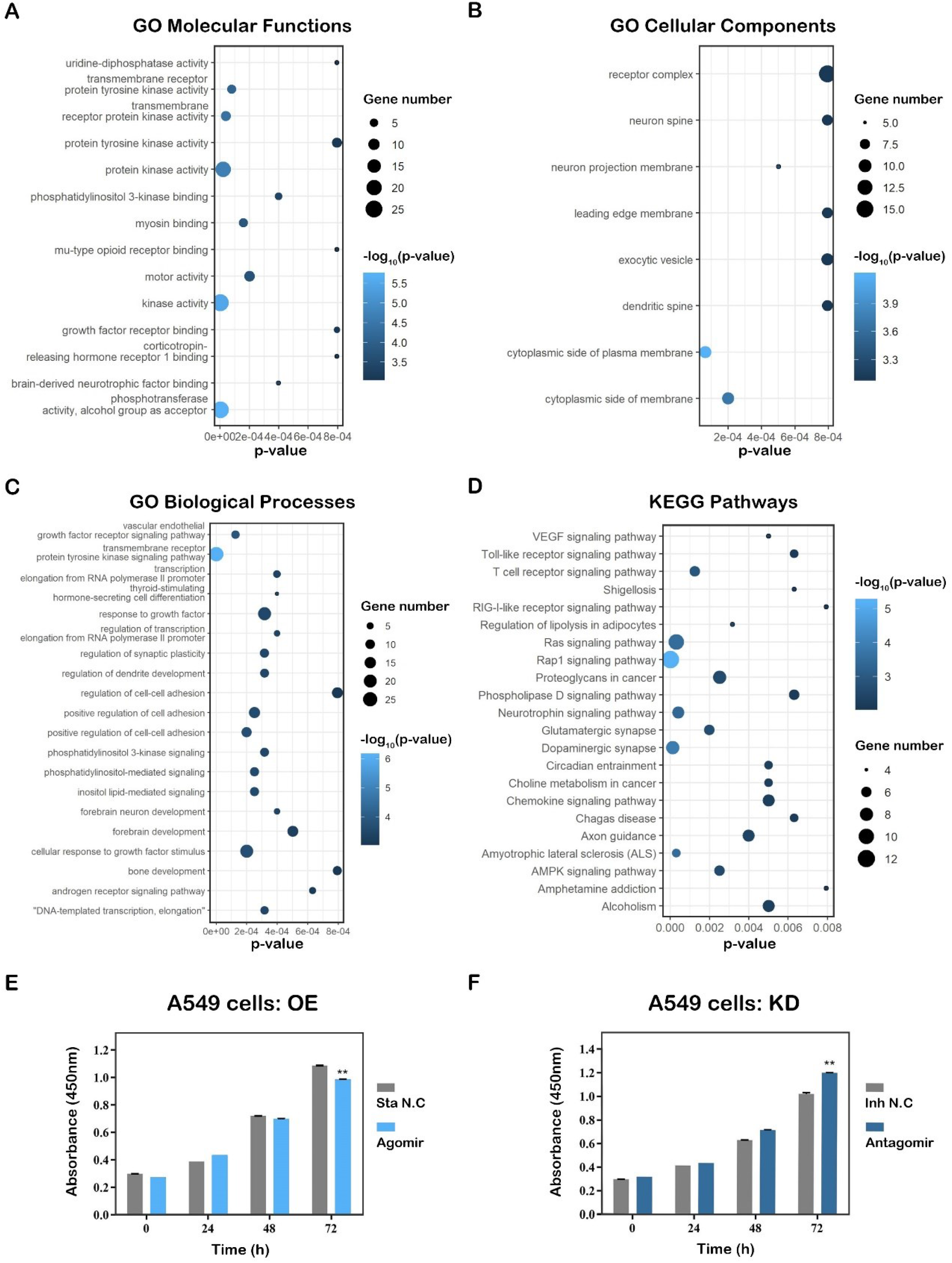
Targets-based functional analysis and validation of hsa-miR-654-5p. Gene ontology (GO) annotation analysis and KEGG pathway enrichment analysis results were performed by Metascape and visualized by R package “ggplot2” based on 275 overlapping genes. Each bubble represents a term, and its size represents the counts of involved genes. Lighter colors of bubbles indicate smaller P values. (A) Enriched terms of GO molecular functions (MFs, p<0.001). (B) Enriched terms of GO biological processes (BPs, p<0.001). (C) Enriched terms of GO cellular compounds (CCs, p<0.001). (D) Enriched terms KEGG pathway (p<0.01). (E and F) A549 and H1299 cells were transfected with miR-654-5p agomir or antagomir respectively. 36 h after transfection, CCK8 cell vitality assay was performed to detect the cell proliferation capacity of miR-654-5p-overexpressing A549 cells (E) and miR-654-5p-knockdown H1299 cells (F). Data from the CCK8 proliferation assay represent the mean±SEM of 3 independently prepared samples. *p<0.05, **p<0.01, ***p<0.001 vs. control.

Hallmark gene sets, oncogenic signatures, Reactome, and KEGG functional sets were also performed by Metascape based on the 275 genes, and the results showed that in the KEGG functional sets, PI3K-Akt signaling and MAPK (p38 and JNK) signaling were enriched (Supplementary Fig. S2, Supplementary Fig. S3 E and F). For Reactome, RTK-related items like signaling by the RTK, VEGF and VEGFA-VEGFR2 pathways were enriched as expected (Supplementary Fig. S2D), further proving the regulatory role of miR-654-5p in cell proliferation. For Hallmarks enrichment, items about interferon response epithelial-mesenchymal transition (EMT) were enriched (Supplementary Fig. S2 A). In oncogenic signature, TGF-β up, STK33 down, ESC v6.5 up, CAMP up and AKT up/mTOR down were most significantly enriched (Supplementary Fig. S2 C). Among these results, TGF-β is a key inducer for EMT of cancer cells. AKT up and mTOR down indicates inhibition of the PI3K-AKT-mTOR pathway, which might lead to cell proliferation inhibition (Supplementary Fig. S2).

In addition, we further performed enrichment analysis based on all high score targets (Score>0.95) predicted by miRWalks 3.0 to avoid missing any key regulatory information resulting from merely taking the intersection of predicted targets into consideration^35^. In GO BPs, except for cell-cell adhesion and MAPK cascade, terms such as ubiquitin-dependent protein catabolic process were most significantly enriched (Supplementary Fig. S2C). As for GO CC, miRNA-related RISC complex, RISC-loading complex and ciliary base were enriched (Supplementary Fig. S2B). In KEGG pathway enrichment, terms such as ubiquitin mediated proteolysis, neurotrophin signaling pathway and cancer-related items such as glioma, hedgehog-signaling pathway and hepatocellular carcinoma were enriched (Supplementary Fig. S2D), which indicated that miR-654-5p might be involved in various aspects of carcinogenesis and tumor progression.

To validate the effect of miR-654-5p on lung cancer cells proliferation *in vitro*, we overexpressed miR-654-5p in A549 cells. 36 hours after transfection, CCK8 cell proliferation assays were subsequently performed to assess the proliferative capacity of the cells. The results showed that the miR-654-5p-overexpression group showed higher proliferative ability (Fig. 3E, p < 0.01) compared to the control group, indicating that upregulation of miR-654-5p inhibits the proliferation of lung cancer cells. To further prove our hypothesis, another lung cancer cells H1299 were transfected with antagomir to knockdown the endogenous abundance of miR-654-5p. Proliferation assays were then performed, and the results showed that decreased miR-654-5p promoted the proliferation capacity (Fig. 3F, p < 0.01) of lung cancer cells. In summary, these results showed miR-654-5p inhibited lung cancer cell proliferation *in vitro*, which are consistent with our previous functional analysis of miR-654-5p.

#### 3.4.2 Abundance-based functional analysis shows that miR-654-5p inhibits lung cancer cell migration

To further study the role of miR-654-5p in lung cancer, pan-cancer normalized LUAD mRNA dataset were divided into three group based on the miR-654-5p of samples (Fig. 4A and B), and differential expression analysis was subsequently performed using package “limma”. Relative to low abundance group, differential expression genes (DEGs) such as MEG3, COL12A1, SPOCK1, IGF2, SFRP2 are significantly elevated, and CA4. FIGF, PGC, GPX2 are markedly decreased in high abundance group (Fig. 4C and D). subsequent functional enrichment analyses showed that these genes were enriched in lung disease, lung small cell cancer, PI3K-Akt-mTOR pathway and the epithelial-mesenchymal transition processes (Fig. 4E).

**Figure 4.**
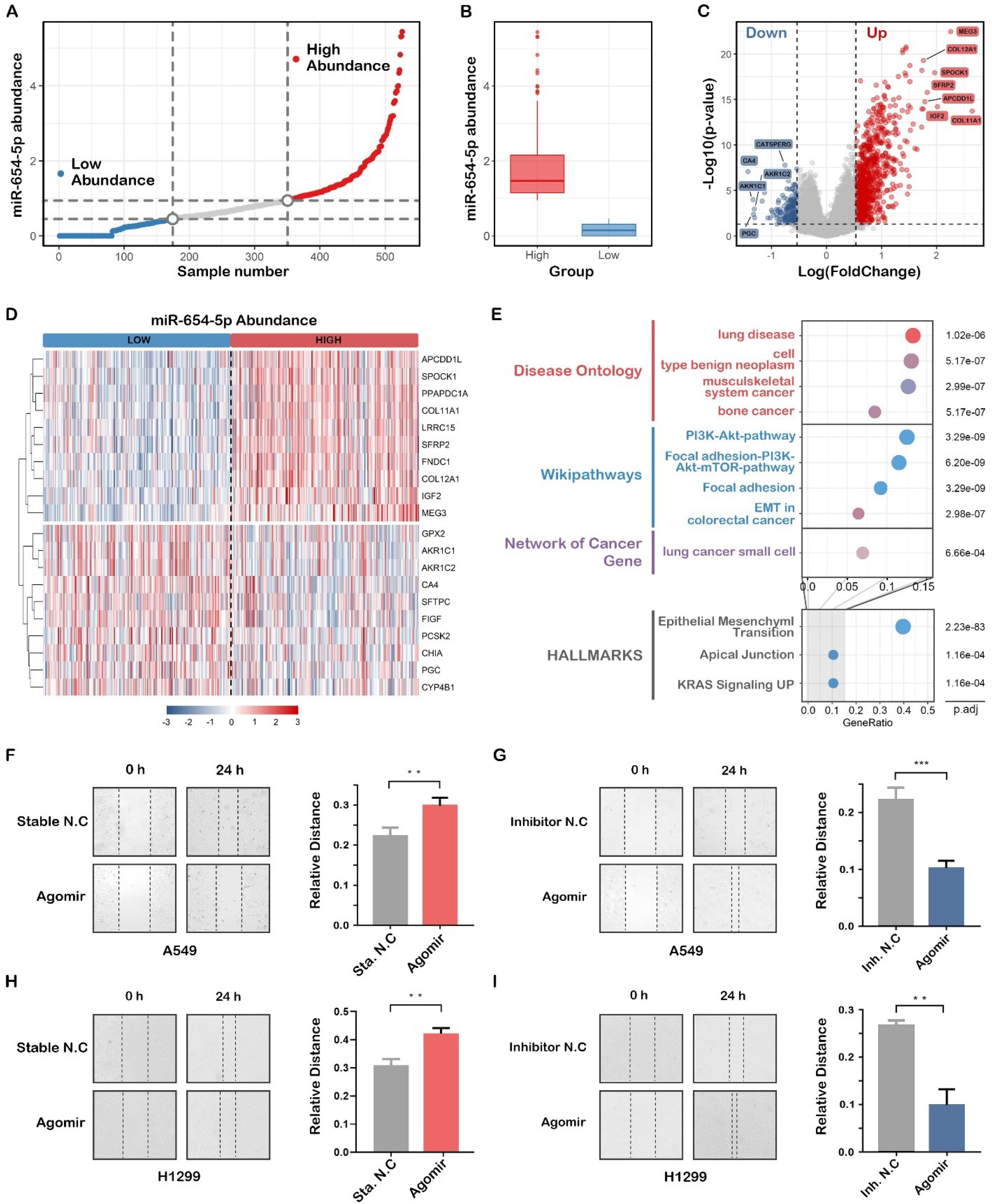
Abundance-based functional analysis of hsa-miR-654-5p. (A) LUAD samples in pan-cancer batch effect normalized mRNA expression dataset was ordered by miR-654-5p abundance and then divided into three groups. (B) The abundance of hsa-miR-654-5p in the high abundance group and the low abundance group. (C) Volcano plot for the differential expression analyses. (D) heatmap for top10 differentially expressed genes. (E) Functional enrichment analysis for differentially expressed genes based on R package “msigdbr” and “clusterProfiler”. (F and G) Representative microscopic images and quantitative results of migratory cells from the miR-654-5p-mimic (agomir)-transfected A549 lung cancer cell group (F) and the miR-654-5p-inhibitor (antagomir)-transfected A549 lung cancer cell group (G) in wound healing assays; (H and I) Representative microscopic images and quantitative results of migratory cells from the miR-654-5p-mimic (agomir)-transfected H1299 lung cancer cell group (H) and the miR-654-5p-inhibitor (antagomir)-transfected H1299 lung cancer cell group (I) in a wound healing assays. Data from the wound healing assay represent the mean±SEM of 3 independently prepared samples. *p<0.05, **p<0.01, ***p<0.001 vs. control.

Taken together, miR-654-5p targets-based enrichment analysis and miR-654-5p abundance-based functional both indicated that hsa-miR-654-5p might be involved in lung cancer cell proliferation, migration and EMT processes.

To prove the effect of miR-654-5p lung cancer cells on migration, wound healing assays were performed to assess the migratory capacity of the cells. The results showed that the miR-654-5p-overexpression group showed higher mobility (Fig. 4F, p < 0.01; Fig. 4H, p < 0.01) compared with the control group, while decreased miR-654-5p promoted migration capacity (Fig. 4G, p < 0.001; Fig. 4I, p < 0.01) of lung cancer cells. In summary, these results showed miR-654-5p inhibited lung cancer cell migration *in vitro*, consistent with our previous analysis.

#### 3.4.3 ssGSEA-based functional analysis shows that miR-654-5p inhibits epithelial-mesenchymal transition

Both targets-based enrichment analysis and abundance-based functional analysis implied that miR-654-5p might regulate epithelial-mesenchymal transition, a key process in lung cancer metastasis. To further reveal the correlation between miR-654-5p abundance and lung cancer cell migration, ssGSEA analysis was subsequently performed. The results showed that both in LUAD and LUSC, miR-654-5p abundance were negatively associated with the EMT score of samples (Fig. 5A and 5C). Compared to high miR-654-5p abundance group, lower hsa-miR-654-5p significantly increased EMT score (Fig. 5B and 5D), indicating a progression of epithelial-mesenchymal transition of lung cancer cells. These results reveal the negative correlation between miR-654-5p abundance and EMT process.

**Figure 5.**
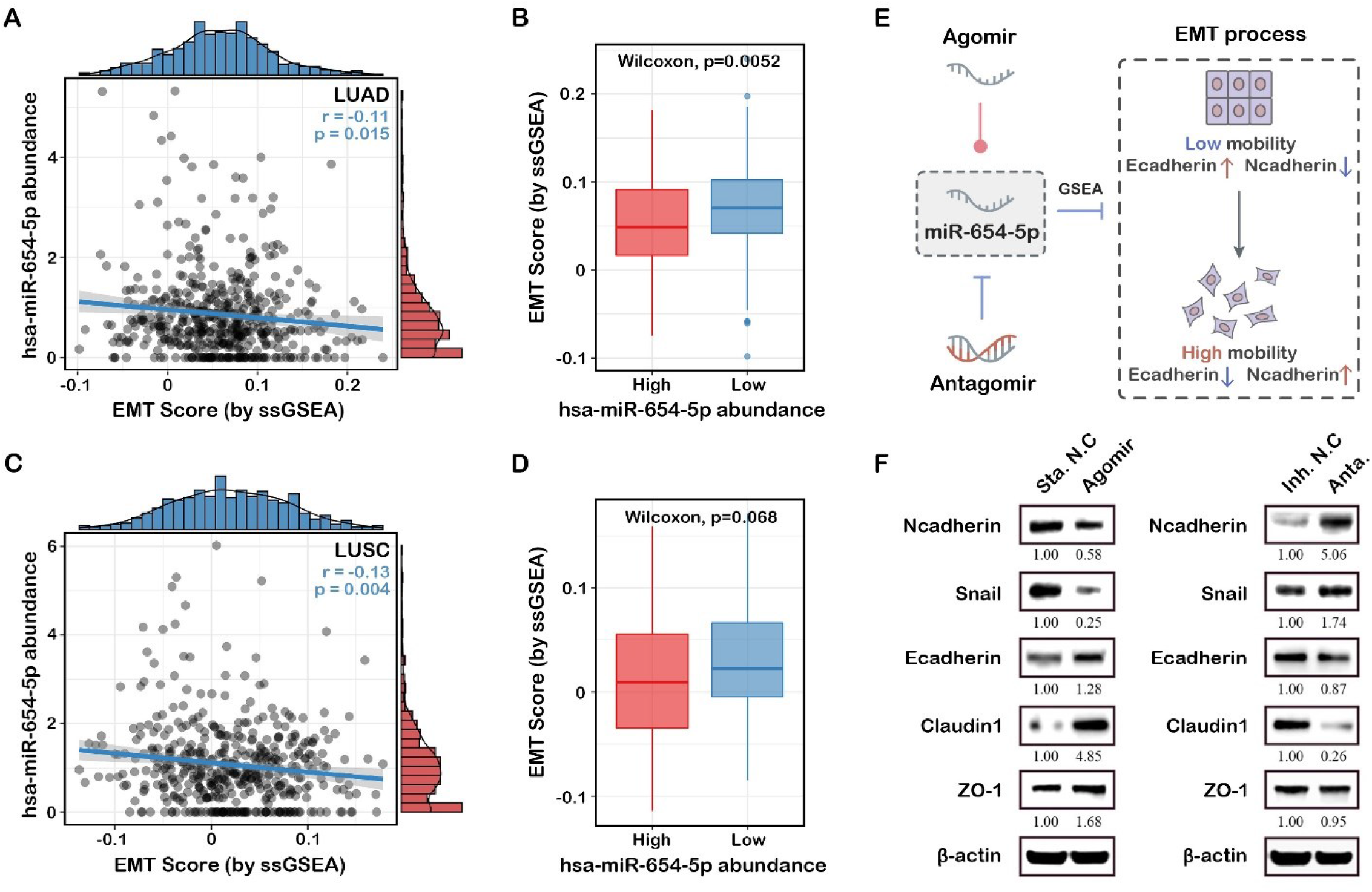
ssGSEA-based functional analysis of hsa-miR-654-5p. The correlation between miR-654-5p and ssGSEA-based EMT process scores were assessed in LUAD(A) and LUSC(C). EMT score in miR-654-5p high abundance group were compared to the low abundance group in LUAD(B) and LUSC(D). (E) Graphical representation of experimental validation of miR-654-5p functions *in vitro*. (F) A549 and H1299 cells were transfected with miR-654-5p agomir or antagomir respectively. 36 h after transfection, the cells were harvested and lysed to extract total proteins, then western blot assays were performed to detect changes in EMT-related markers (epithelial status hallmarks: E-cadherin, ZO-1 and claudin-1; mesenchymal status hallmarks: Ncadherin and Snail).

EMT can be characterized via the expression level of EMT-related signatures, which included epithelial status markers such as E-cadherin, ZO-1 and mesenchymal status markers such as vimentin, N-cadherin and Snail (Fig 5E)^36-38^. To confirm that miR-654-5p can regulate EMT as we analyzed, we upregulated and downregulated miR-654-5p in lung cancer A549 cells by agomir and antagomir, respectively. The results showed that the evaluated miR-654-5p could downregulate the expression of N-cadherin (to 0.58-fold, Fig. 5F) and Snail (to 0.25-fold, Fig. 5F), while enhancing the expression of E-cadherin (by 1.28-fold, Fig. 5F), ZO-1 (to 1.68-fold, Fig. 5F) and Claudin-1 (to 4.85-fold, Fig. 5F), indicating that overexpression of miR-654-5p inhibited the EMT process in lung adenocarcinoma cells. In miR-654-5p-knockdown cells, the results indicated the downregulation of epithelial markers (to 87% for E-cadherin, 95% for ZO-1, and 26% for Claudin-1) and the upregulation of mesenchymal markers (by 406% for N-cadherin and 74% for Snail; Fig. 5B).

Previous enrichment analysis indicated that cell adhesion was a potential function of miR-654-5p, while EMT is known to decrease cell-cell adhesion and give cancer cells migratory and invasive capacity^36,37,39,40^. We reasonably concluded that miR-654-5p might regulate cell migration and invasion via downregulating EMT.

### 3.5 Construction of PPI network and identification of hub target genes

To elucidate the potential interactions among the 275 overlapping genes, a protein-protein network was constructed utilizing the STRING database. A network including 98 nodes and 120 edges was constructed (Fig. 6A). These 98 genes might be crucial in miR-654-5p-regulated cellular processes especially those highly connected to others such as PIK3R1 and RHOCA. By combining these 98 genes with those differentially expressed in lung adenocarcinoma (LUAD) and those highly connected (Degree >2), we found that there are 11 genes in the intersection including HIST2H2BE, RGS4, and RAB10. these predicted targets of miR-654-5p might play pivotal roles in lung adenocarcinoma (Fig. S5).

**Figure 6.**
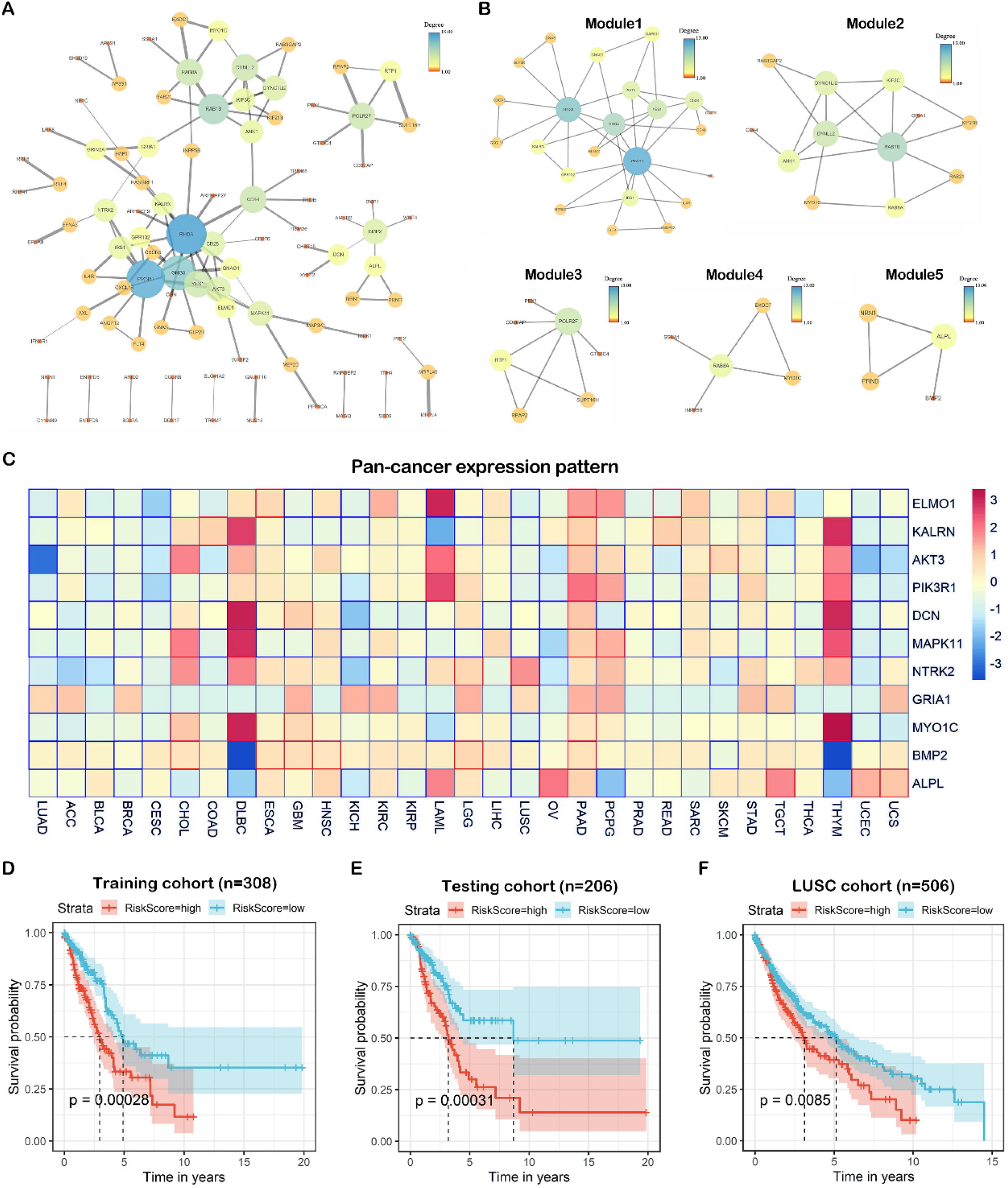
The comprehensive analysis of target genes. (A) A network with 98 nodes and 120 edges was constructed by the STRING database for the overlapping 275 genes predicted by five promising miRNA-targets prediction tools with a high confidence (interaction score>0.700). Disconnected nodes were removed. Edges connecting nodes symbolized the interaction between two genes and the wider edges indicated a higher combined score between two nodes. The color and size of the nodes indicates the degree of the nodes. The network was visualized by Cytoscape 3.7.1 software. (B) built-in app MCODE was utilized to find modules in the whole network. Five modules were identified. (C) Pan-cancer differential expression heatmap was normalized and plotted by R package “pheatmap” based on TCGA cancer samples. The color in each block represents the median expression ratio of a gene (tumor/normal). Differentially expressed genes are framed in red (upregulated) or blue (downregulated), a P-value<0.05 was considered to indicate a statistically significant difference. (D) the LUAD samples were divided into two groups based on risk score, the Kaplan-Meier survival analysis was performed to assess the survival rate of two group in training cohort (n=308). (E) A testing cohort generated via randomly sampling of TCGA LUAD data set, the overall survival analysis was then performed. (F) The risk score model was applied to an independent LUSC cohort

To find the potential interconnected regions in this network, MCODE was utilized, and five networks were clustered (Fig. 6B). In these networks, AKT3, RAB1B, RTF1, EXOC7, and ALPL were identified as seeds, indicating these genes might be the promising pivotal genes in miR-654-5p-regulated biological processes.

On the other hand, 275 targets of miR-654-5p with differential expression and differential survival rate in various cancers were also screen in order to provide more specific suggestions for miR-654-5p-related functional research in various cancers (Table S1).

### 3.6 Pan-cancer expression and prognostic value of hub genes

To understand the roles of the 11 hub targets in various cancers, we analyzed the expression patterns of the hub genes in cancer and normal tissue in the TCGA database (Fig. S6A), all hub genes were significantly downregulated in lung cancer compared to normal. Pan-cancer expression heatmap showed that these hub genes exhibited distinct expression patterns in different cancer. For example, most hub genes were downregulated in LUAD, BLCA and BRCA, while most genes were upregulated in PAAD, indicating the roles of miR-654-5p/targets/axes were different in various cancers. Interestingly, a similar expression pattern of hub genes could also be found. For instance, the hub genes expression pattern in THYM and DLBC, showing the unification of correlations between miR-654-5p/targets/axes and a certain type of cancer (Fig 6C).

To assess the prognostic value of the hub genes, Kaplan-Meier survival analysis was performed based on the 11 hub genes selected by the PPI network in the LUAD TCGA database, in order to acquire a promising judgement as to whether these hub genes can be prognostic markers. The results showed that in LUAD, high expression of MAPK11 and MYO1C (Fig. S6B) was negatively related to patients’ survival rate, while high expression of AKT3, ALPL, DCN, ELMO1, GRIA1, KALRN, NTRK2 and PIK3R1 was correlated with a better prognosis (Fig. S6B). These genes can be potential prognostic markers for lung cancer. Furthermore, the survival rate heatmap including OS (Fig. S6C) of these hub genes was also plotted based on the TCGA database, which might be used as a reference for prognostic markers.

To further identify the potential prognostic value of the hub genes, TCGA LUAD samples were randomly divided into training cohort (n=308) and testing cohort (n=206), univariate cox regression and multivariate cox regression were then performed to assess the prognostic value of hub genes. Following stepwise regression, a three-gene risk scoring model were constructed:

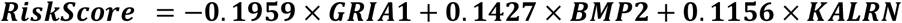

Both in training cohort and testing cohort, we found that high risk score was associated with poor prognosis (Fig. 6D and E), the similar results could also be observed in an independent LUSC validation cohort. The ROC curve showed that the AUC values of the model were 0.677 at 5 years of OS (Fig. S7A). to further test the prognostic value of this model, we apply the Riskscore to pan-cancer mRNA dataset, results showed that the AUC values of model are 0.8, 0.78, 0.77, 0.74, 0.64 in PRAD, UVM, USC, PAAD, TGCT respectively (Fig. S7B), indicating the scalability of the hub genes-based Riskscore model.

## 4 DISCUSSION

Currently, multiple researchers have confirmed that miRNAs are involved in the mechanisms of most biological processes including excessive growth, resistance to apoptosis, angiogenesis, invasion, and metastasis^28,40-42^. Accordingly, aberrant miRNA expression patterns are considered to be a sign of a variety of diseases such as cancer, suggesting the expression of miRNAs could be used as diagnostic, prognostic and predictive biomarkers^43^. The function of some miRNAs in cancer has been clarified and divided into oncogenic miRNAs (oncomiRs) or anti-oncogenic miRNAs (tumor suppressor miRs), For example, Let-7a, an identified tumor suppressor, has been shown to inhibit the proliferation of lung cancer cells by regulating HMGA2^44^ and myc^45^. In the present study, miR-654-5p abundance in tissues were found to be correlated to the progression and survival rate in multiple cancers such as lung cancer, which were consistent with previous reports^14^. These findings highlight the importance of miR-654-5p-related research. However, due to the complexity of mammalian cells, the miRNA regulatory network seems to be much more complex than a simple miRNA-target-phenotype regulation pattern. Identifying the target set of miR-654-5p and its potential functions would therefore provide a larger map reference for the mechanism underlying miR-654-5p-regulated biological processes. To address the issue, we designed a workflow integrating *in silico* identification, comprehensive functional analysis, clinical value assessment of miR-654-5p hub targets and functions. As a proof of concept, we also validated miR-654-5p-RNF8 axis *in vitro*. Our results not only identified a new functional target for hsa-miR-654-5p, but also demonstrated our effective target prediction strategy of miR-654-5p.

To obtain a better and common understanding of regulatory pattern and functions of miR-654-5p at a cellular level, three kinds of data-driven bioinformatic methods including target-based, abundance-based and ssGSEA-based functional analysis was performed. Our results first implied that miR-654-5p might be a crucial regulator of cell growth and proliferation. We then proved that miR-654-5p inhibited cell proliferative capacity, while downmodulation of miR-654-5p promoted cell proliferation in lung cancer. Our finding was consistent with many previous reports^15,16,46^. As a matter of fact, targets-based functional analysis is a general method that its results can be applied to different certain cancer types, which gives us a clue to what to do next. Meanwhile, subsequent abundance-based functional analysis and many other bioinformatic analysis can be further integrated with results from target-based analysis to further demonstrate the correlation between miRNA and a certain phenotype like what we did in this article.

Multiple functional analysis results showed that miR-654-5p are negatively associated with lung cancer EMT processes, which is a pivotal process of cancer metastasis known to decrease cell-cell adhesion and give cancer cells migratory and invasive capacity^36,37,39,40^. We reasonably suggest that miR-654-5p might regulate cell adhesion via regulation of the EMT process. In OSCC, this mechanism had been demonstrated by Lu^16^. However, the relation between hsa-miR-654-5p and EMT in other cancers type are yet to be discovered. Based on these previous findings and *in vitro* experiments, we demonstrated the miR-654-5p indeed inhibited the EMT process and cell-cell adhesion. In addition, wound healing and Transwell assays proved the inhibitory function of miR-654-5p on cell migration in LUAD, which confirmed our functional analyses result as an *in vitro* proof of concept. However, the correlation between miR-654-5p with cancer cell metastasis remained to be discovered.

Meanwhile, we also noticed that miR-654-5p might play important roles in many common biological processes. For example, neuron-related items such as dendrite and forebrain neuron development, dopaminergic synapse were enriched, indicating that miR-645-5p might be pivotal in neuron-related functions. Interestingly, a study focused on the correlation between miRNA and neurodevelopmental disorders revealed that miR-654-5p is commonly deregulated in autism spectrum disorders (ASD)^47^, indicating an association between miR-654-5p and neurodevelopmental disease like ASD, which needs further study. Bone development, ubiquitin-related items such as ubiquitin protein ligase activity and ubiquitin-mediated proteolysis, and cancers such as glioma and hepatocellular carcinoma were enriched. The relationship of miR-654-5p to these enriched terms needs to be studied further.

For pathways, we found MAPK-related signaling, PI3K/AKT pathway, T cell receptor (TCR) signaling, and Ras signaling were all enriched in multiple enrichment analysis, and these pathways could be given priority when we further study the regulatory pattern of miR-654-5p. Through literature searching, we found that some of our analyses have already been proven, such as the regulatory function of miR-654-5p on the MAPK pathway in OSCC^16^ and osteogenic differentiation^48^. Thus, our functional analysis provides a framework for studying miR-654-5p biological functions.

We found that most hub genes were downregulated in LUAD, BLCA and BRCA, and upregulated in PAAD. Of note, the hub genes’ expression patterns in THYM and DLBC were similar, which needs to be explored in the future. Meanwhile, we showed that all hub genes were significantly correlated with patients’ survival and these genes could be utilized as prognostic markers for LUAD. Additionally, the results showed that using GRIA1, BMP2 and KALRN together as a signature was of great value for the prognosis of LUAD. However, the risk scoring based on stepwise modeling of these genes were not that satisfactory (AUC=0.677 at 5 years), combining them with clinical factors might enhance their capability as a prognostic prediction model.

To conclude, our research utilized a close-loop experiment integrating bioinformatic analysis and *in vitro* experimental validation to explore the function of the poorly studied miR-654-5p, aiming to provide a larger map reference for the functions and regulation network of miR-654-5p. In addition, we, for the first time, experimentally validated the direct regulatory effect of miR-654-5p on RNF8 as proof of concept of our target prediction. We also predicted, revealed, and then validated the regulation functions of miR-645-5p on the lung cancer cell EMT process, cell proliferation and migration capacity for the first time, which not only proved our bioinformatic functional analyses but also revealed multiple biological functions that miR-654-5p might regulate.

## 7 Author Contributions

CL, LM and LZ designed the experiments.CL performed bioinformatic analysis the experiments, miR-654-5p-RNF8 *in vitro* functional validation and wrote the manuscript. LM and JK performed and analyzed the *in vitro* data. CZ and JM contributed to the construction of dual-luciferase vector. XQ and XM contributed to functional analyses. CL, LM, LZ and SX edited the manuscript. LZ contributed to funding acquisition. **All authors read and approved the final manuscript**.

## 9 Acknowledgments

None.

## 10 Data Availability Statement

The data that support the findings of this study are openly available in GEO database (https://www.ncbi.nlm.nih.gov/geo/) and TCGA database (https://xenabrowser.net/)

---

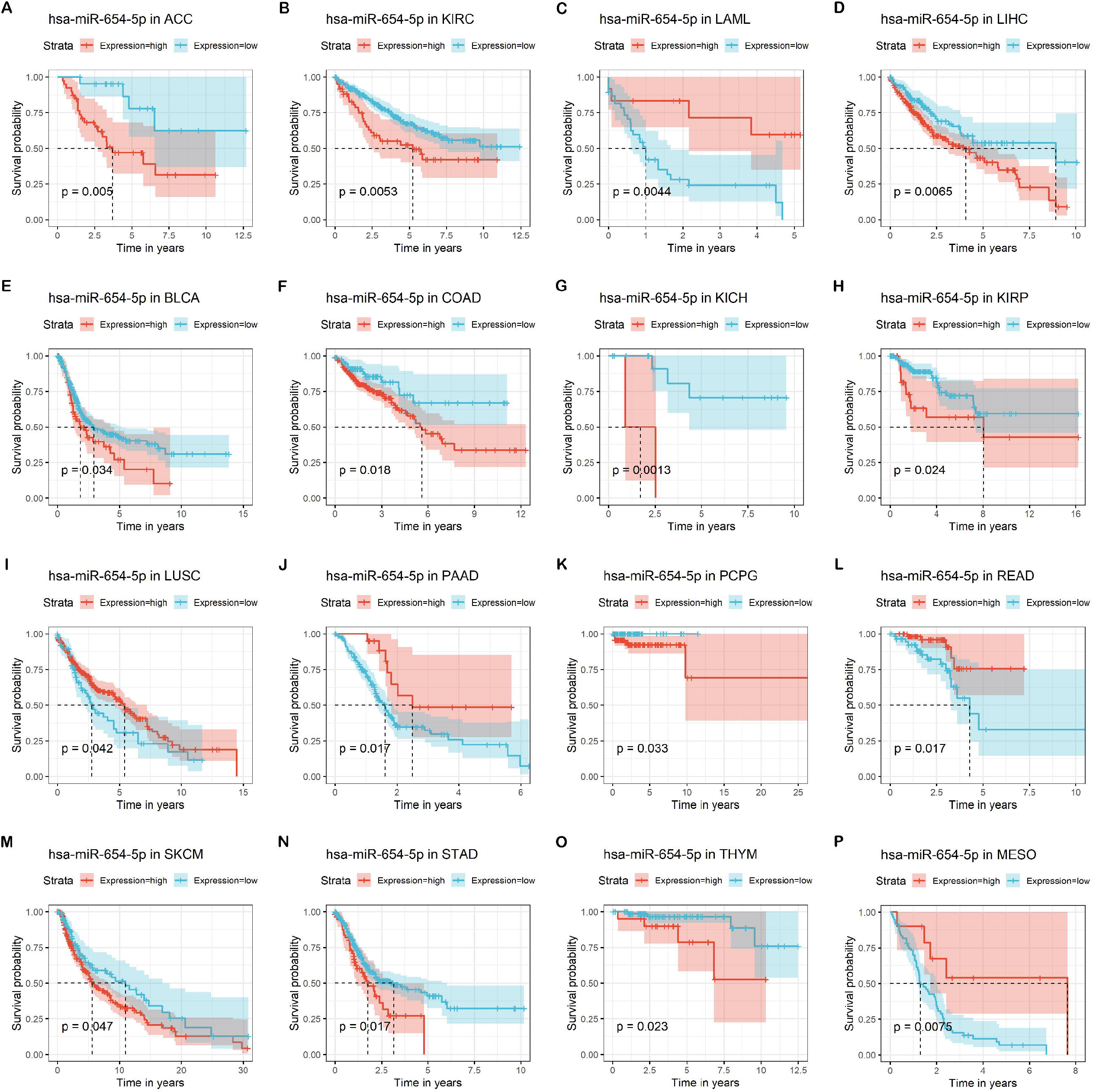

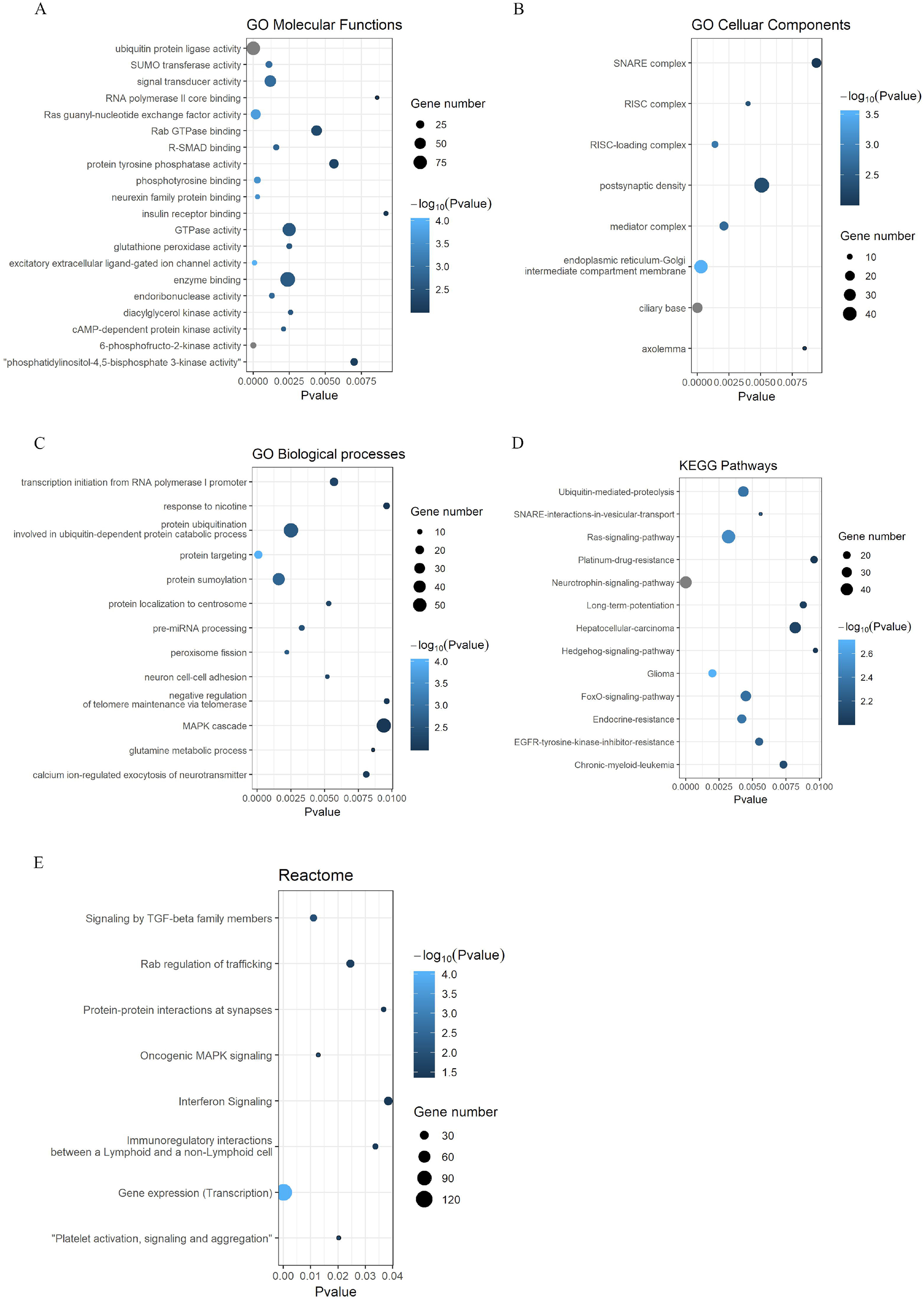

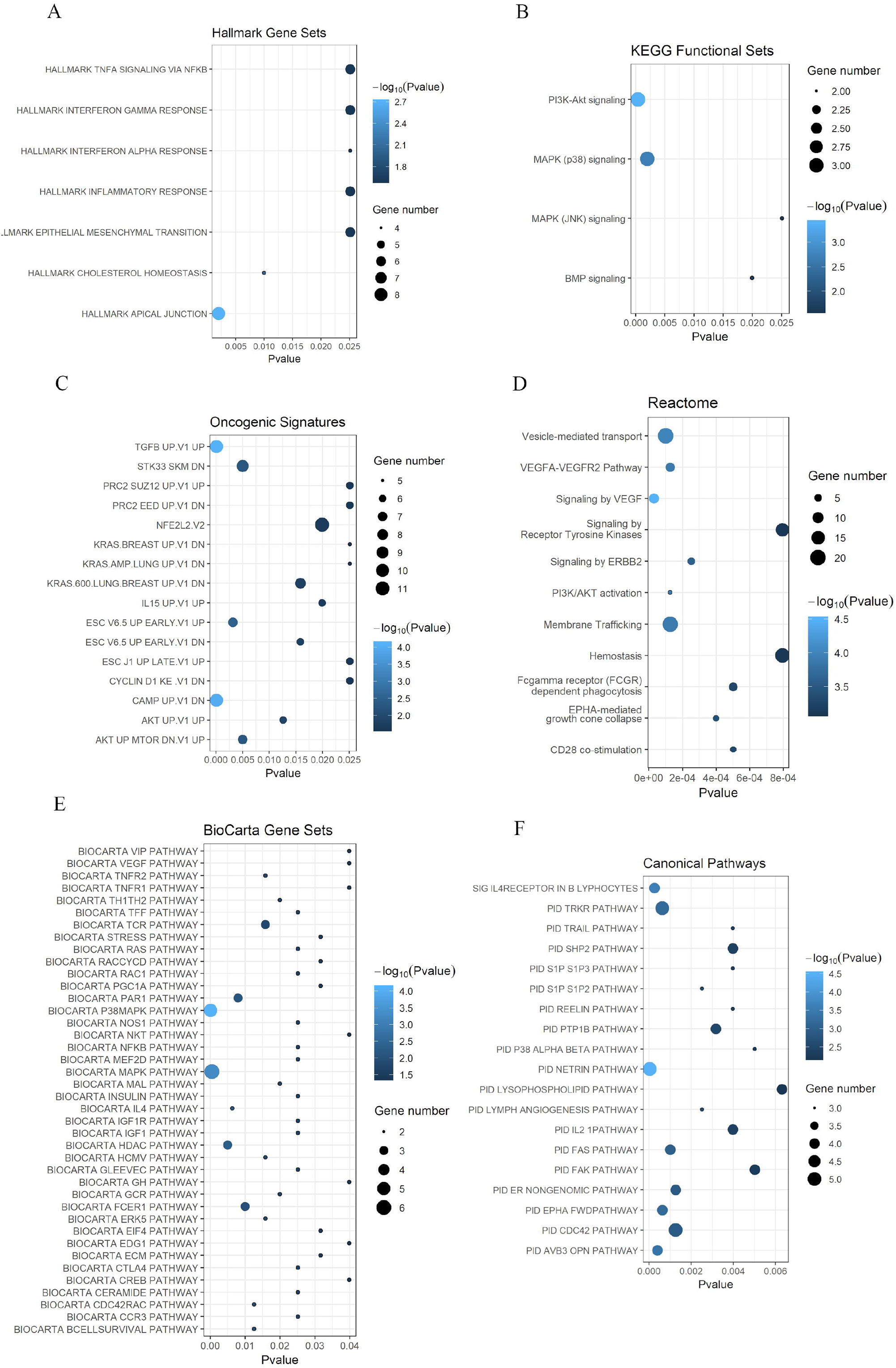

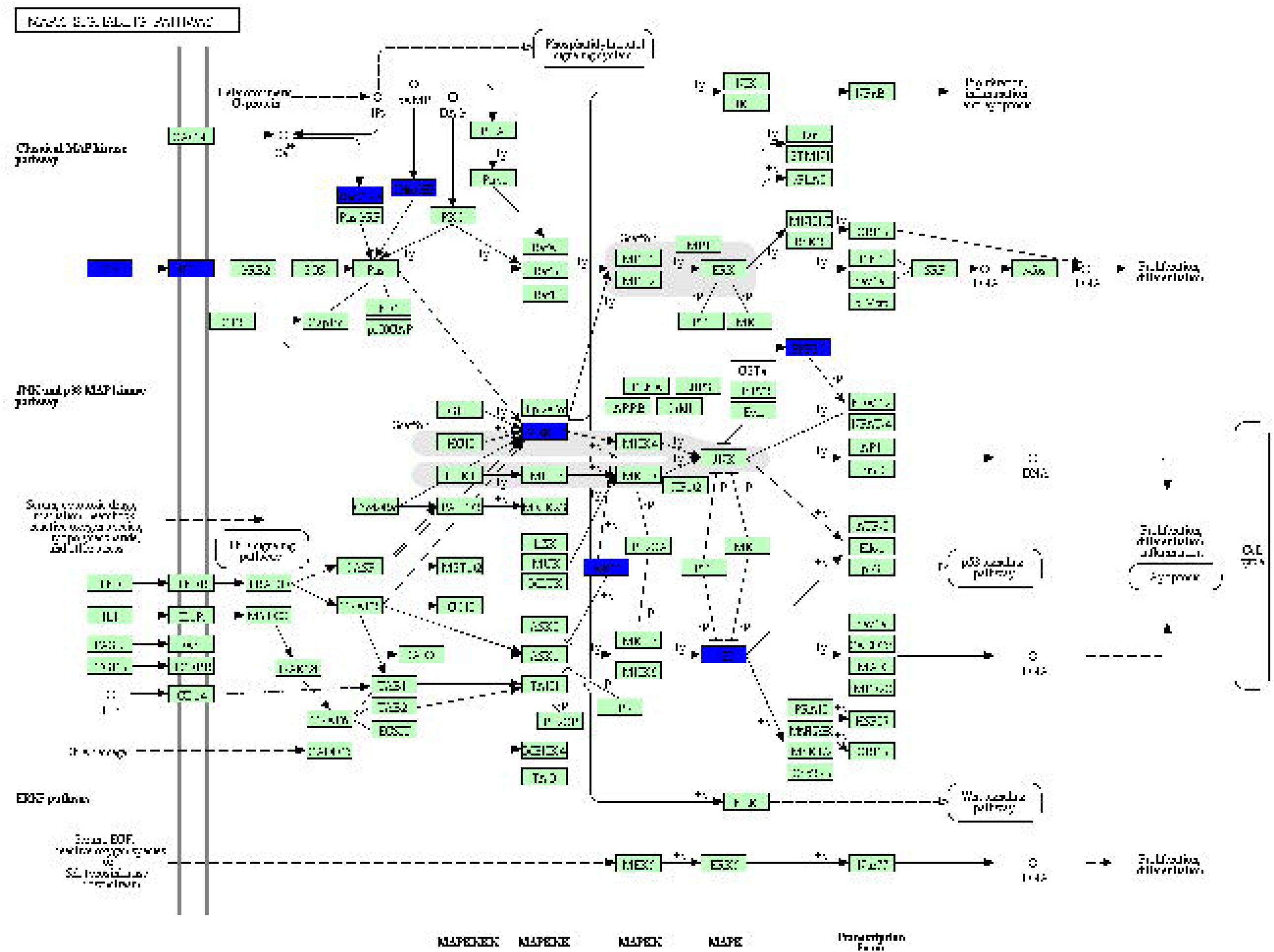

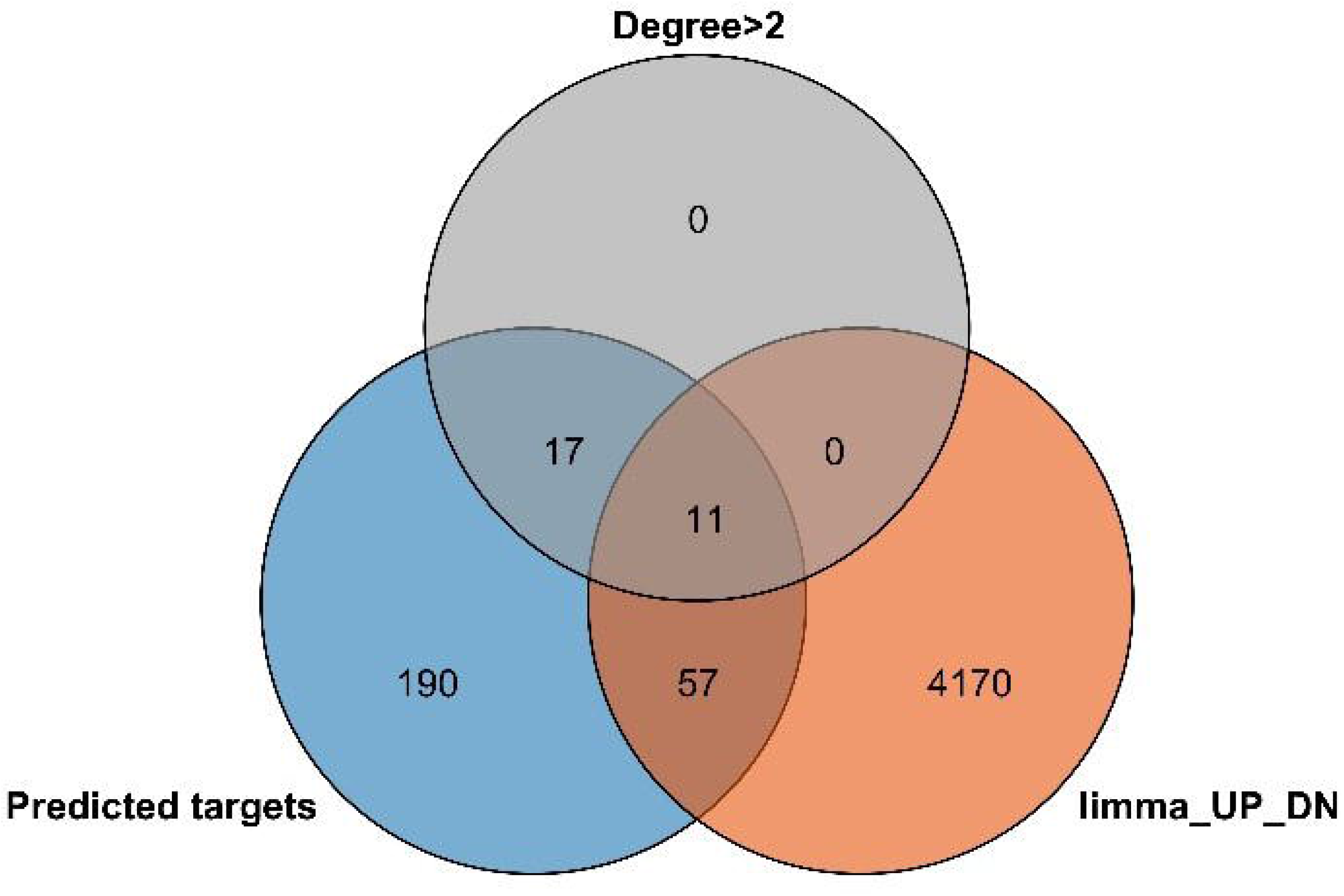

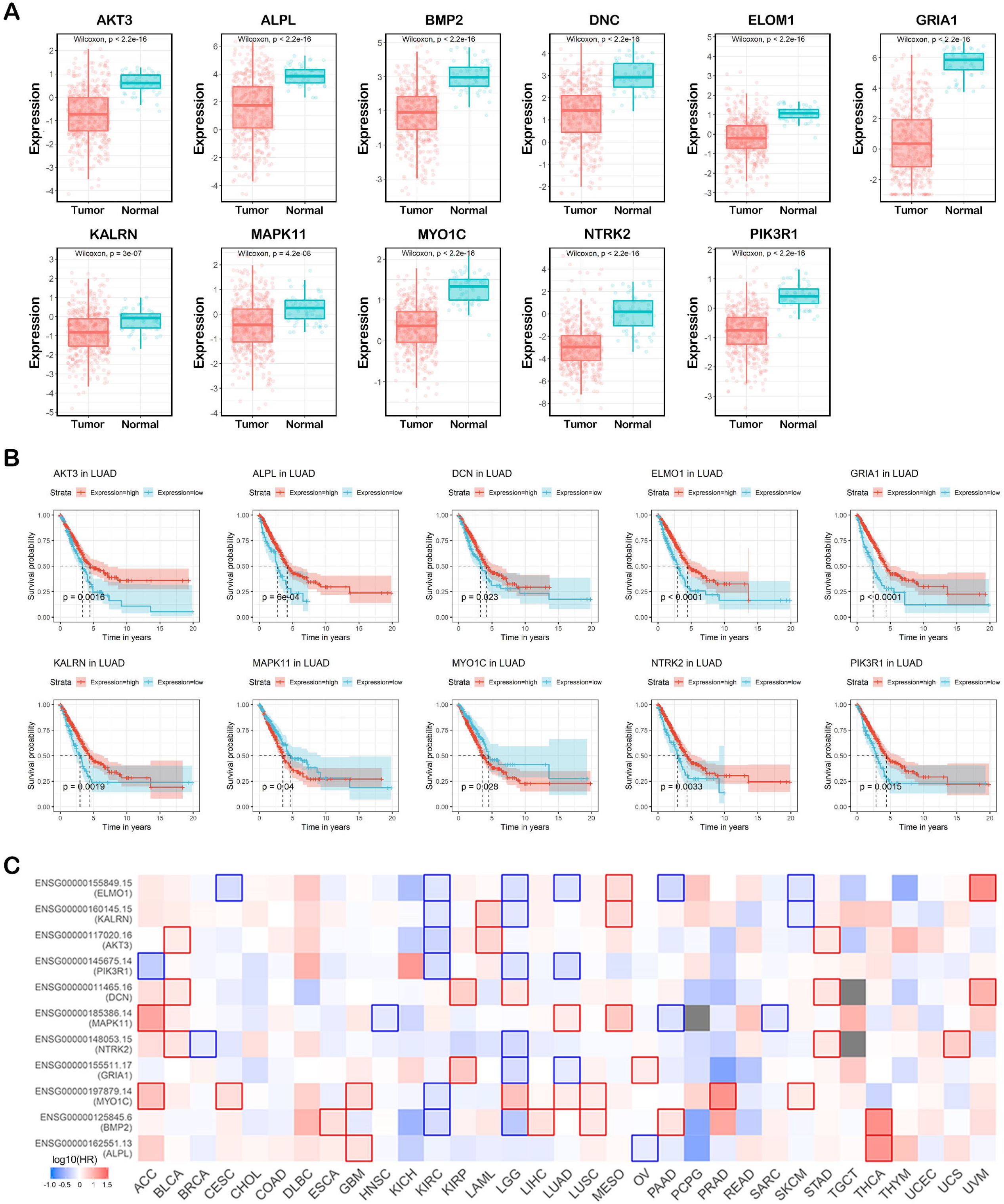

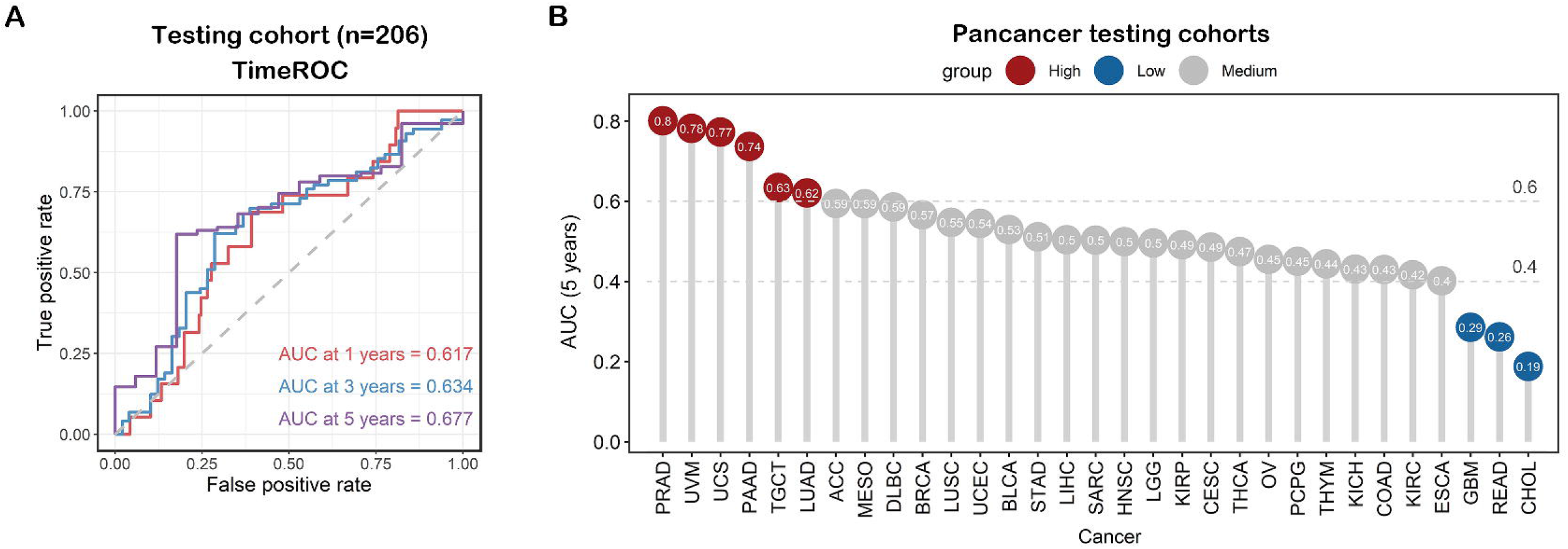

