## Supplementary material for "Comprehensive analysis of hsa-miR-654-5p’s tumor-suppressing functions": Fig. S

### Supplementary Materials

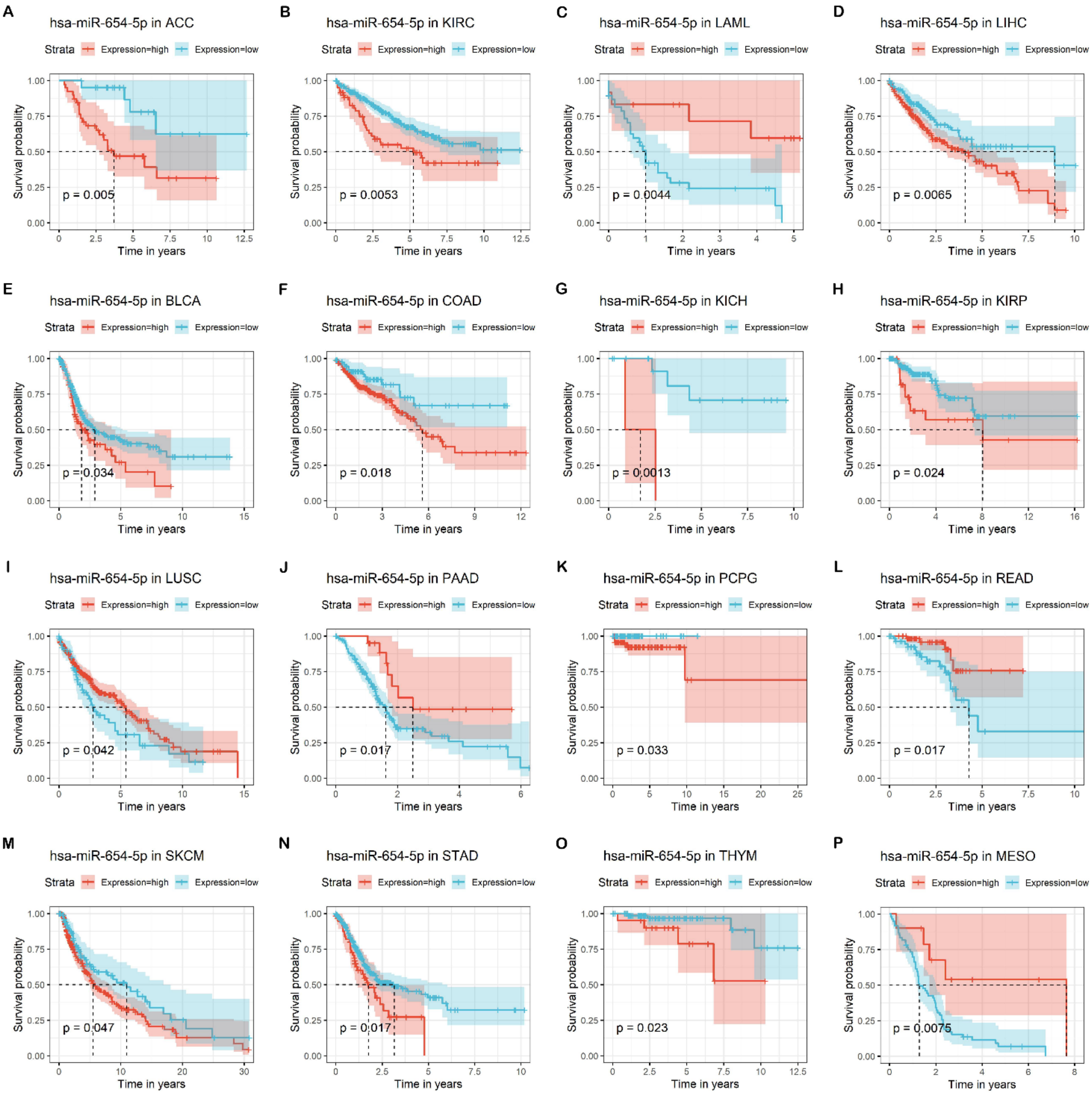


**Figure S1. Survival analysis of hsa-miR-654-5p in the pan-cancer dataset.**

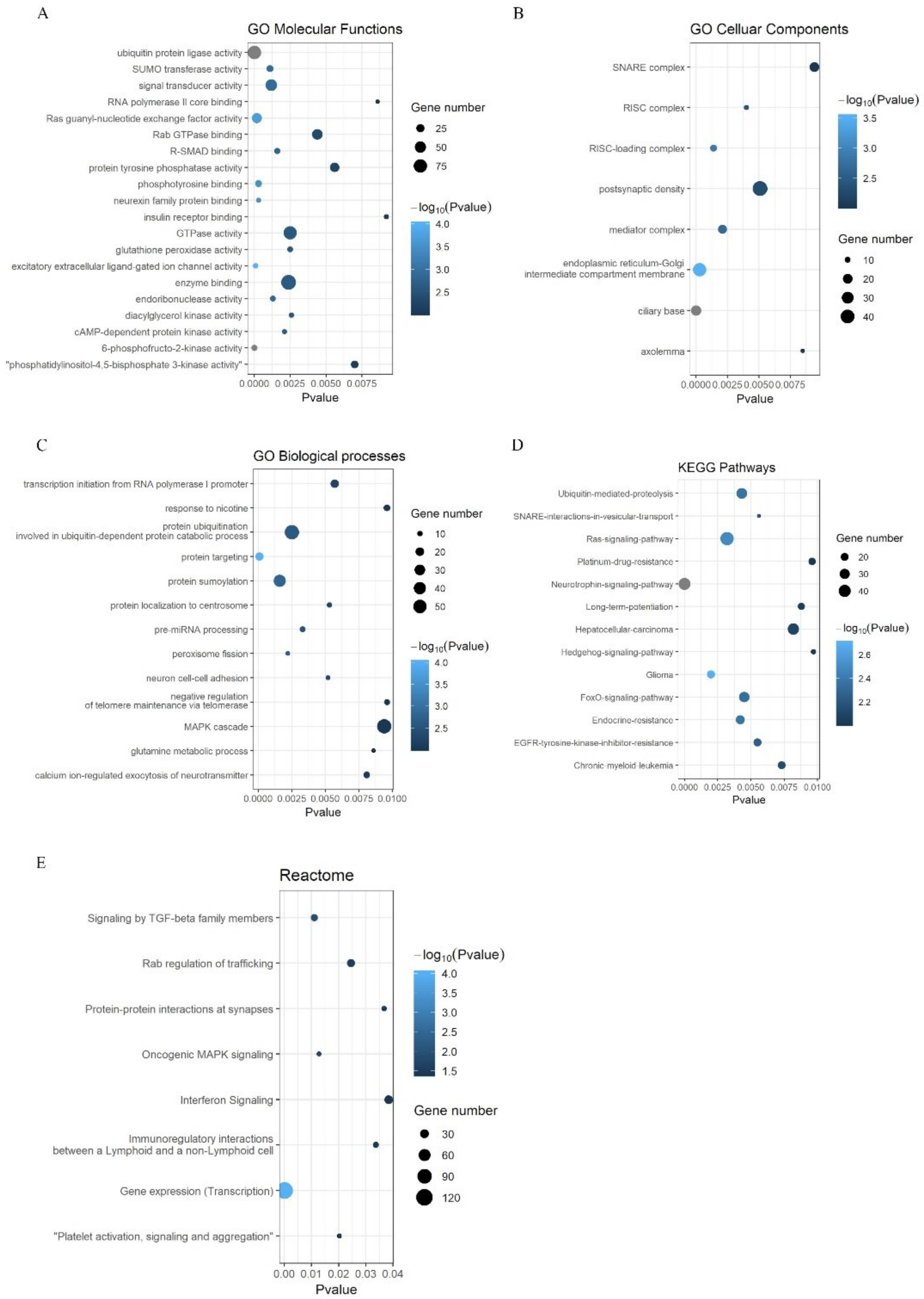


**Figure S2. Gene ontology and KEGG pathway enrichment analysis of predicted targets of miR-654-5p by miRWalks3.0.** the predicted genes (score>0.95) of miR-654-5p by new version 3.0 of miRWalks were all chosen to perform GO annotation and KEGG pathway enrichment analysis results to . Each bubble represents an enriched term, its size represents the counts of involved genes. Lighter colors indicate smaller P values. (A) Enriched terms of GO molecular functions. (B) Enriched terms of GO cellular compounds. (C) Enriched terms of GO biological processes. (D) Enriched terms of KEGG pathway. (E) Enriched terms of Reactome.

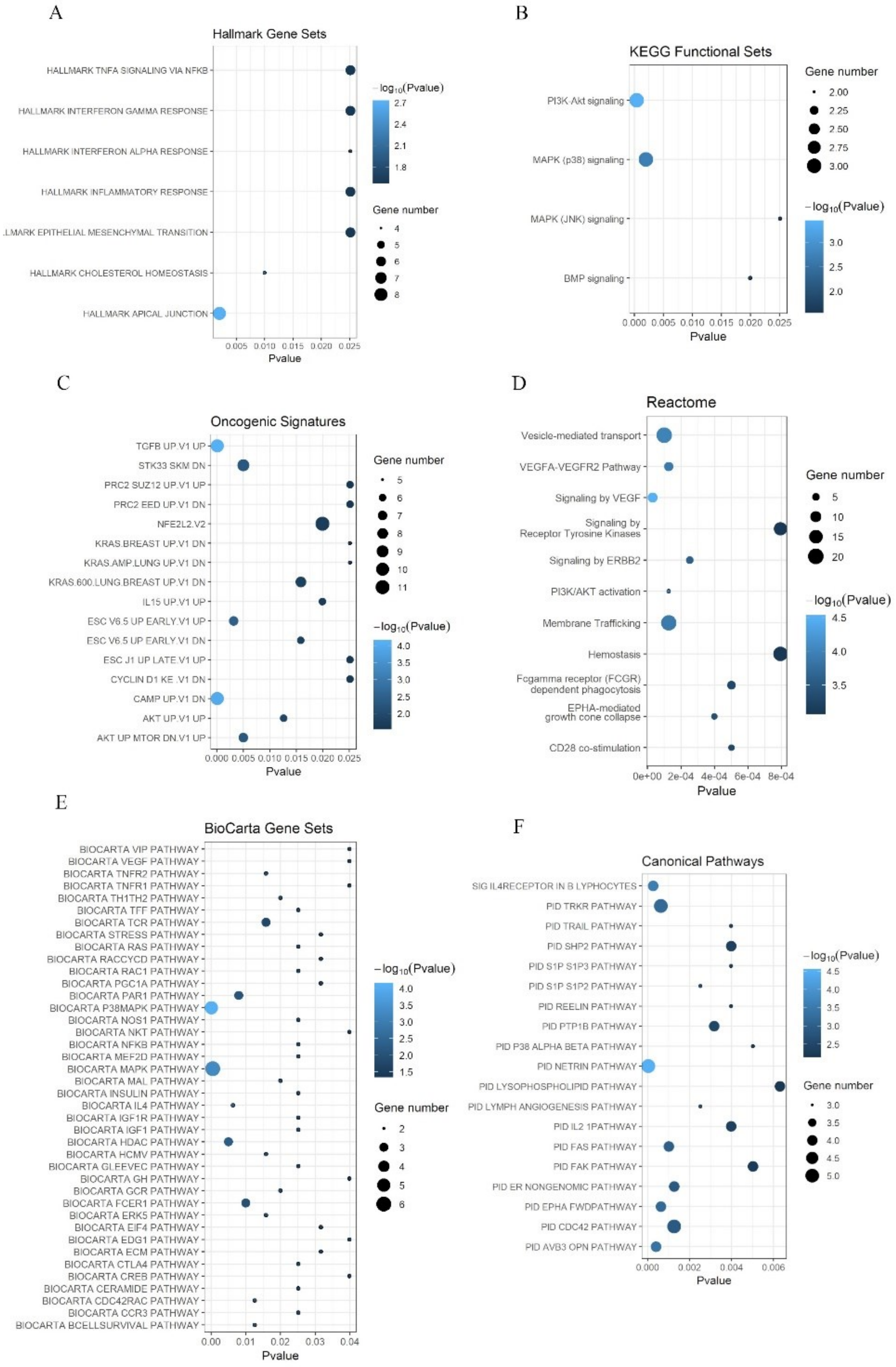


**Figure S3. enrichment analysis of 275 overlapping predicted targets of miR-622.** the 275 predicted genes of miR-654-5p were chosen to perform Hallmark, KEGG functional sets, Oncogenic signatures, Reactome, BioCarta and Canonical pathway enrichment analysis. Each bubble represents a term, and its size reprensent the counts of involved genes. Lighter colors indicate smaller P values. (A) enriched items for hallmark gene sets ($p<0.05$). (B) enriched items for KEGG functional sets ($p<0.05$). (C) enriched items for Oncogenic signatures ($p<0.05$). (D) enriched items for Reactome ($p<0.001$). (E) enriched items for BioCarta Gene Sets ($p<0.05$). (F) enriched items for Canonical Pathways ($p<0.01$)[1, 2].

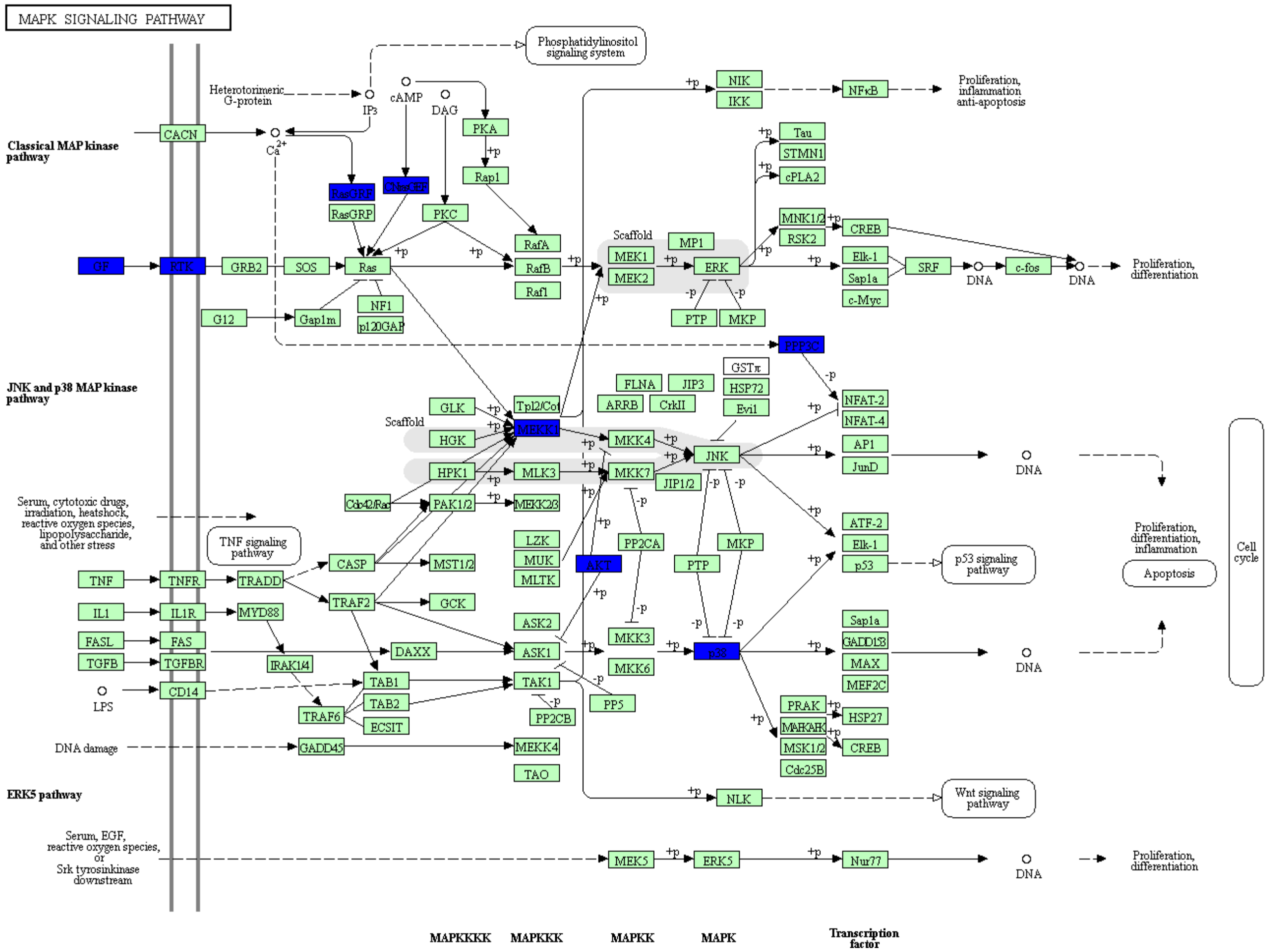


**Figure S4. The inhibitory influence of miR-654-5p on MAKP pathway in *Homo sapiens*.** the 275 predicted genes of miR-654-5p were mapped to KEGG pathway. Genes regulated by miR-654-5p were colored in blue.

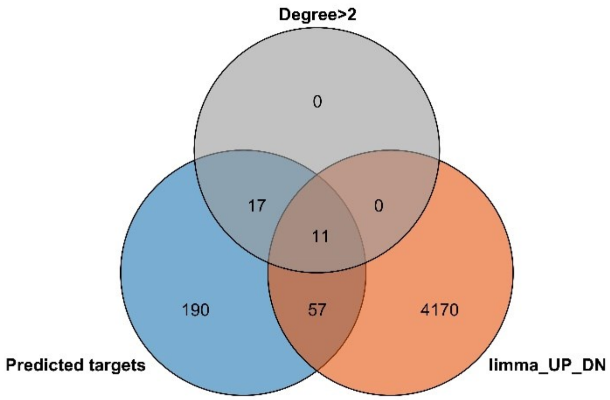


**Figure S5. The hub genes of miR-654-5p in LUAD.** 275 predicted genes were screened by the criterion of high connection in PPI network (Degree>2) and differential expressing (by R package limma with corrected p-value<0.05) in TCGA LUAD database[3].

**Table S1. The identification of key targets of miR-654-5p in various cancers.**

| ACC | BLCA | BRCA | CESC | COAD | DLBC | ESCA |
|---|---|---|---|---|---|---|
| ASF1B | ATP8B2 | - | PEAR1 | - | POLR2F | ATF3<br>GNL3L<br>ANGPT2 |
| **GBM** | **HNSC** | **KICH** | **KIRC** | **KIRP** | **LAML** | **LGG** |
| MYO1C<br>TMEM150A | - | - | - | NRN1<br>MFAP4 | RNF8<br>MRPL49<br>ASF1B | MAP3K1<br>PBX3 |
| **LIHC** | **LUAD** | **LUSC** | **OV** | **PAAD** | **PRAD** | **READ** |
| - | PPT2<br>PIK3R1 | CADM3<br>BMP2<br>FLT4<br>ASF1B | ALPL<br>GALNT10<br>PEAR1 | ATP8B2<br>AP3S1<br>ASF1B | - | ZNF493<br>NICN1 |
| **SKCM** | **STAD** | **TGCT** | **THCA** | **THYM** | **UCEC** | **UCS** |
| - | EFNA3<br>HIF3A<br>AFF3<br>UBE2QL1<br>ASF1B | PEAR1<br>ZNF385B | - | SLC7A6<br>HDAC7 | PEAR1 | SYT13 |

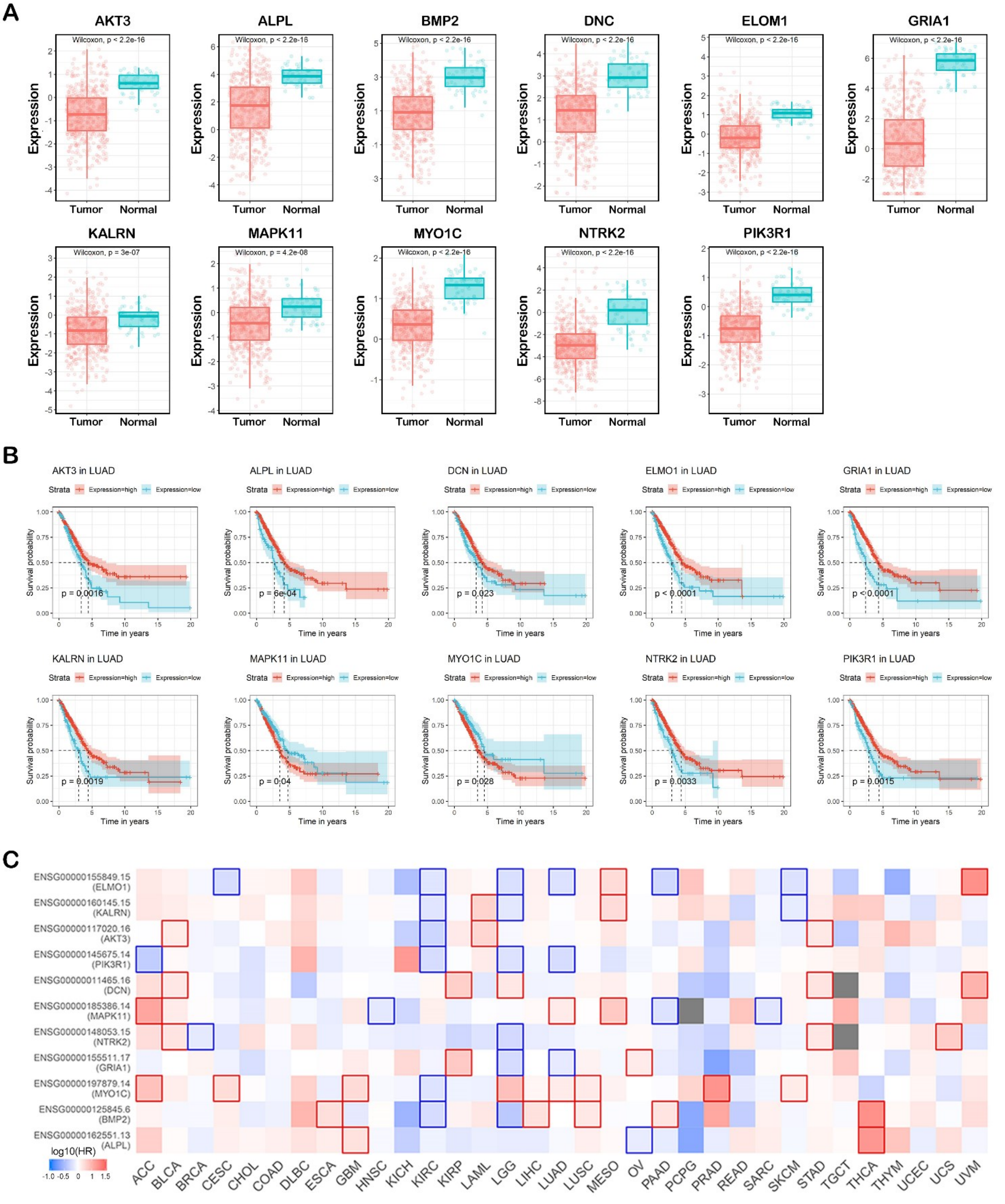


**Figure S6. The expression and prognostic analysis of 11 hub targets in cancers** (A) The differential expression analyses of hub genes in LUAD were based on pan-cancer batch effect normalized TCGA samples, Wilcoxon test was used to compare the mean between the cancer group and the normal tissue group. (B) The OS rate correlated with 11 hub targets by Kaplan-Meier survival analysis based on the TCGA LUAD database. A log rank $p < 0.05$

was considered to indicate a statistically significant difference. Those with $p>0.05$ are not shown. (C) The heatmap of the pan-cancer OS rate of 11 hub targets by Kaplan-Meier survival analysis based on TCGA samples by GEPIA. A log rank $p <0.05$ was considered to indicate a statistically significant difference and are framed in red (positively correlated) or blue (negatively correlated).

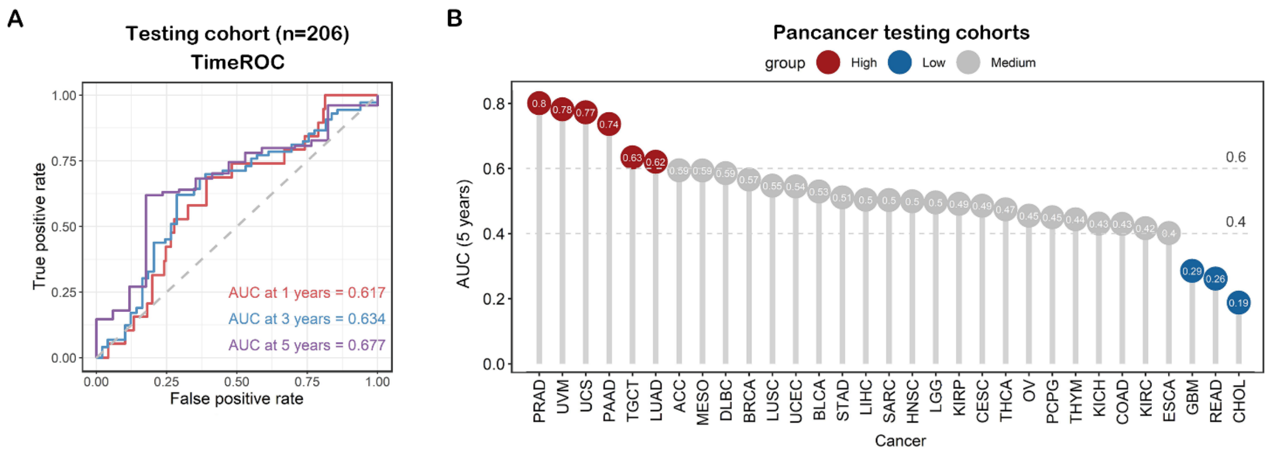


**Figure S7.** The performance of the risk score model. (A) ROC curve showing the model performance in testing cohort. (B) The risk score model was applied to Pan-cancer mRNA dataset.
